# Dorsal forerunner cells couple epiboly movements to zebrafish notochord extension

**DOI:** 10.1101/2025.10.24.684459

**Authors:** Christopher Small, Margot Kossmann Williams

**Author notes:** Department of Molecular Genetics, The Ohio State University, Columbus OH.

## Abstract

Convergence and extension (C&E) cell movements promote anteroposterior axis extension and narrow both neuroectodermal and mesodermal tissues during gastrulation. Mediolateral cell intercalation is largely responsible for this morphogenetic process in vertebrates, but evidence suggests that additional cell-extrinsic forces generated by surrounding tissues contribute to the shaping of many structures. In zebrafish, for example, mechanical forces generated by the anteriorly migrating prechordal plate cooperate with tissue-autonomous cell intercalations to promote extension of the notochord. Here we propose a novel model for notochord morphogenesis by which mechanical epiboly forces within the enveloping layer are transmitted to the posterior end of the notochord via a cluster of dorsal forerunner cells (DFCs) that physically links the two. We found that scattering or absence of DFCs caused by loss of *crb2a* or *sox32*, respectively, reduces notochord C&E and exacerbates axis extension defects in planar cell polarity signaling-deficient embryos. Using an automated image segmentation and cell shape analysis pipeline, we show that cells within the posterior notochord fail to properly elongate when DFCs are scattered or absent. Finally, we demonstrate that loss of *crb2a* and *sox32* fails to disrupt C&E of zebrafish embryonic explants in which no epiboly occurs and all extension is driven by cell-intrinsic behaviors. Together, these findings support a model in which DFCs facilitate mechanical coupling of the enveloping layer to the posterior notochord during epiboly to ensure its robust morphogenesis during gastrulation.

## INTRODUCTION

During gastrulation, the newly formed germ layers - mesoderm, endoderm and ectoderm - are shaped into a nascent body plan by a suite of morphogenetic cell behaviors. Among these is convergence & extension (C&E), the concomitant mediolateral (ML) narrowing and anteroposterior (AP) elongation of tissues that extends the head-tail axis ^1,2^ and enables neural tube closure ^3–6^. Planar polarized cell behaviors including ML cell elongation, alignment, junctional shrinking, and protrusive activity drive intercalations to rearrange cells into longer and narrower arrays ^7^. The polarization of these cell behaviors in vertebrate embryos requires planar cell polarity (PCP) signaling, which orients cells and their behaviors with respect to the AP axis ^8^. While the critical role of PCP signaling in vertebrate gastrulation morphogenesis is well established ^9–12^, additional cues regulating C&E independently of PCP remain incompletely understood.

Cell-extrinsic forces (i.e. those generated by neighboring tissues) also contribute to axial extension. Mesoderm internalization in *Drosophila* gastrulae generates mechanical forces that promote passive germ band extension ^13^, and anterior migration of the prechordal plate and head mesendoderm of zebrafish and amphibian embryos, respectively, contributes to C&E by pulling on the chordamesoderm to which they are attached ^14–17^. Embryonic landmarks such as tissue boundaries also promote C&E, as contact with natural or induced boundaries triggers polarized C&E cell behaviors in the chordamesoderm ^18–20^ in a PCP-independent fashion ^21,22^. Morphogen signaling gradients also promote C&E independent of PCP, including graded JAK/STAT signaling in the *Drosophila* hindgut ^23^ and Nodal signaling in vertebrates ^24–27^. Nodal is best known for specifying mesoderm and endoderm ^28–31^, but studies suggest its role in AP axis extension is at least partially independent of mesoderm formation ^24–26^. Although a few genes activated downstream of Nodal were found to regulate morphogenetic cell behaviors ^32^, it remains poorly understood how Nodal activity promotes C&E during gastrulation.

Because Nodal signaling largely functions to alter gene expression through the transcriptional co-regulator Smad2 ^33^, we sought to identify Nodal transcriptional targets with roles in C&E morphogenesis. By intersecting our own published datasets of genes differentially expressed between wild-type (WT) and Nodal signaling deficient maternal-zygotic (MZ)*oep*-/-zebrafish gastrulae ^26^ and those induced by Nodal activation within otherwise naïve zebrafish explants that undergo C&E ^34^, we identified *crumbs2a* (*crb2a*) as a candidate morphogenetic effector downstream of Nodal. Crumbs2 is a key component of the Crumbs apical polarity complex, which is essential for formation and function of apical cell surfaces and associated cellular structures such as cilia ^35–40^. Crumbs proteins interact with other apical proteins such as atypical protein kinase C (aPKC) and PARs 3 and 6 ^38,40,41^, FERM (4.1, Ezrin, Radixin, Moesin) domain proteins like Yurt and Moesin ^42–44^, and members of the Hippo signaling pathway ^43,45,46^. In multiple species and developmental contexts, Crumbs exhibits anisotropic localization along cell interfaces and/or tissue boundaries, where it becomes mutually exclusive with Myosin ^47–49^. This is reminiscent of the polarized Myosin localization underlying *Drosophila* germ band elongation ^50,51^, providing a potential mechanism by which Crumbs2 could instruct vertebrate C&E cell behaviors. Indeed, *Crb2*-/-mouse gastrulae have severely shortened and widened AP axes, consistent with C&E defects ^49^. Zebrafish embryos mutant for *crb2a* (also known as *oko meduzy* (*ome*)) exhibit failed lamination of retinal cell types, reduced retinal pigmentation, deflated brain ventricles, and failed heart trabeculation ^35,37,52–54^. However, a role for *crb2a* during zebrafish gastrulation was never examined.

Here, we report that loss of *crb2a* significantly reduces C&E specifically in the developing notochord of zebrafish gastrulae. Unlike its broad expression in the mouse epiblast ^49^, however, we find that zebrafish *crb2a* expression during gastrulation is largely limited to the dorsal forerunner cells (DFCs), a small cell cluster on the embryo’s dorsal side that moves with the enveloping layer (EVL) as it undergoes epiboly ^55–57^. DFCs maintain contact with the posterior end of the developing notochord and give rise to the Kupffer’s vesicle upon completion of epiboly ^56–58^. We find that DFCs fail to properly cluster in *crb2a* mutant embryos, and that loss of DFCs phenocopies *crb2a*-/-C&E defects. Quantitative live imaging reveals that failure of DFC clustering in *crb2a* mutants disrupts ML cell polarization in the notochord underlying its C&E. Finally, we show that loss of *crb2a* or DFCs fails to disrupt C&E of zebrafish embryonic explants in which no epiboly occurs and all extension is driven by cell-intrinsic intercalation behaviors. Together, these findings support a model in which the EVL-DFC-notochord nexus couples epiboly movements to enhance C&E of the notochord, providing a new example of how cell-intrinsic and -extrinsic forces cooperate to ensure robustness of a morphogenetic process.

## RESULTS

### Loss of *crb2a* disrupts AP axis extension and notochord C&E

Previous work from our lab identified Nodal signaling as an essential regulator of C&E during zebrafish gastrulation ^26,34^. Evidence suggests that in addition to its role in mesoderm specification, Nodal functions cell autonomously and independently of PCP signaling to promote C&E ^24,26^, but the downstream effectors responsible are unknown. To identify candidate Nodal-dependent regulators of C&E, we examined our own published datasets for genes whose expression is reduced/absent in Nodal signaling-deficient MZ*oep*-/-gastrulae ^26^ and induced by Nodal activation within extending zebrafish explants ^34^. From these intersected datasets, we selected *crb2a* as a promising candidate due to its ability to direct subcellular Myosin localization ^47–49^ and the reduced C&E observed in *Crb2*-/-mouse gastrulae ^49^.

To test if loss of *crb2a* affects C&E in zebrafish, we first performed multi-guide CRISPR targeting ^59^ to disrupt *crb2a* function in injected F0 embryos (hereafter “crispants”). Our cocktail of three single guide (sg)RNAs (**Figure S1A**) efficiently recapitulated *crb2a/ome* mutant phenotypes, including loss of pigment in the retina and heart edema at 48 hours post fertilization (hpf) ^35,52–54^, and loss of Crumbs2a protein expression (**Figure S1C-F**). These embryos had shorter AP axes at day 2, and loss of *crb2a* exacerbated the shortened axes of *vangl2* crispants, in which impaired PCP signaling causes C&E defects ^9,60^ (**Figure S1B-C**). To determine if short axis phenotypes at day 2 reflected C&E defects during gastrulation, we performed morphometric analyses of *crb2a, vangl2,* and *crb2a*+*vangl2* double crispants at the end of gastrulation (tailbud stage, 10 hpf) using whole mount in situ hybridization (WISH) for markers of the neural plate (*dlx3b)*, notochord (*tbxta*), rhombomere 3 (*egr2b*), and prechordal plate (*ctslb*) (**Figure 1A**). *vangl2* crispants recapitulated the wider neural plates and notochords and shortened AP axes of *vangl2* mutants, validating our multi-guide targeting approach to detect C&E defects. *crb2a* crispants exhibited modestly but significantly wider notochords and shorter AP axes, but their neural plates were unaffected (**Figure 1C-E**, left panels). Loss of *crb2a* also exacerbated C&E defects in *vangl2* crispants, but only in the tissues affected in *crb2a* crispants (i.e. the notochord), suggesting that its role is likely independent of PCP signaling.

To ensure that these C&E defects were specific to loss of *crb2a* and not an artifact of multi-guide targeting, we used CRISPR to generate a stable *crb2a* mutant line. While attempting to create a full-locus deletion using two sgRNAs targeting the 5’ and 3’ ends of the gene simultaneously, we isolated a line (designated *bcm122*) with a 1 bp deletion and 4 bp insertion in the last exon of *crb2a.* This caused a frame shift in the critical C-terminal cytoplasmic tail ^42^ (**Figure S1A**) that, when homozygous, perfectly phenocopied *crb2a/ome* mutants ^52,53^ (**Figure S1D**). Crb2a protein was not detected by immunofluorescence in *bcm122* homozygous mutants (**Figure S1G-H)**, demonstrating it is a strong loss-of-function allele. Morphometric analysis of these *crb2a^bcm1^*^22^*^/bcm12^*^2^ (hereafter *crb2a*-/-) stable mutant embryos at 10 hpf (**Figure 1B**) revealed the same phenotype detected in crispants: shorter AP axes and wider notochords, but unaffected neural plates (**Figure 1C-E**, right panels). Together, these results demonstrate that Crb2a promotes C&E specifically within the developing notochord during gastrulation, likely in parallel with PCP signaling.

**Figure 1.**
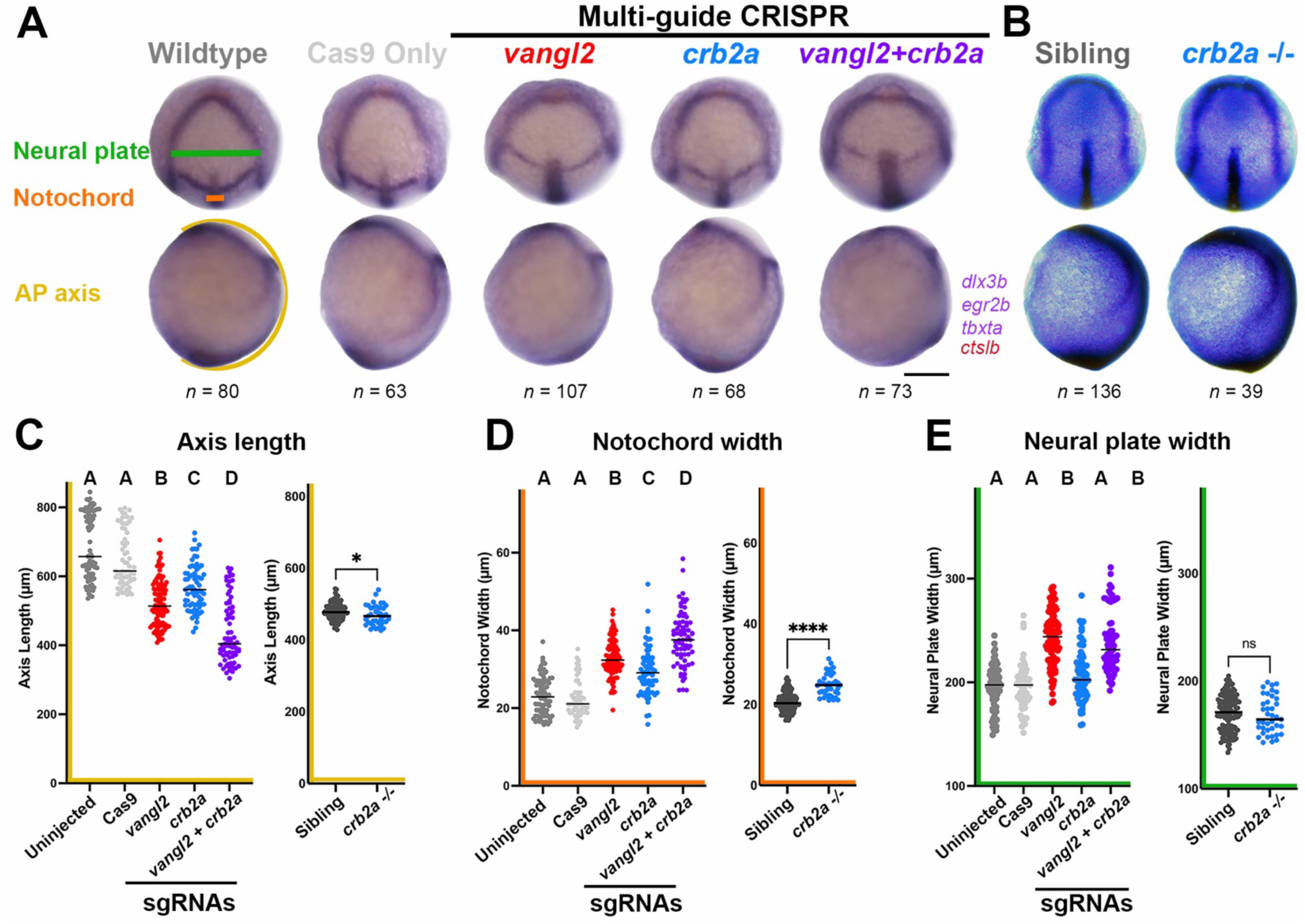
Embryos lacking *crb2a* have shortened body axes and wider notochords, but neural plate width is unaffected. **A-B**) Representative images of WISH for *dlx3b*, *egr2b*, *tbxta*, and *cts1b* in tailbud stage (10 hpf) embryos of the conditions indicated. n = number of embryos measured from three independent trials. **C-E**) Morphometric analyses of axis length (A), notochord width (B), and neural plate width (C) in crispant (left panel) and stable mutant (right panel) embryos shown in A-B. Each dot represents a single embryo, black bars are median values. Crispant data were analyzed with the Kolmogorov-Smirnov test, stable mutants were analyzed with the Mann-Whitney test. Letters denote statistical differences (P < 0.05) between treatment groups, * = P < 0.05, **** = P < 0.0001. See also Figure S1 in Supplemental materials.

### *crb2a* is expressed in the Dorsal Forerunner Cells during gastrulation

Our RNA-seq data indicated that *crb2a* is expressed in zebrafish embryos during gastrulation, but its expression patterns at these stages have not been reported. We therefore examined its expression by WISH from blastomere to early somitogenesis stages (2 – 12 hpf) (**Figure S2**). We first detected *crb2a* expression at shield stage (6 hpf) within a small population of cells at the margin (**Figure S2D**). This expression domain was apparent throughout epiboly stages (**Figure 2A-B**, **Figure S2D-F**) in WT embryos but was not detected in MZ*oep*-/-embryos (**Figure 2C-D**), consistent with its dependence on Nodal signaling at gastrulation stages. Cells expressing *crb2a* at these stages also expressed *sox32* (**Figure 2E-F**) which, combined with their location at the margin, led us to identify them as dorsal forerunner cells (DFCs).

DFCs are a small, transient, and highly specialized cluster of cells that require Nodal for their specification ^57,61^ but, unlike most mesoderm and endoderm cells, fail to internalize at the embryonic margin ^56^. These cells arise from marginal enveloping layer (EVL) cells which divide to produce one EVL cell and one deep cell ^55,62^, some of which become DFCs on the embryo’s dorsal side. DFCs maintain apical attachments to the EVL as they are towed vegetally during epiboly ^55,57^, during which they cluster medially at the dorsal midline in an FGF-^63^ and Eph-Ephrin-dependent ^64^ manner. Immunofluorescence staining with tyramide signal amplification (TSA) revealed that at 75% epiboly, Crb2a protein localized to the membranes of DFCs (**Figure 2I-J**). Although additional background staining is observed with this method, this membrane signal is absent from neighboring deep cells (**Figure 2J**) and from controls without primary antibody (**Figure 2K-L**). Upon yolk plug closure, DFCs condense into a rosette within the tailbud and inflate to form the Kupffer’s vesicle (KV), the zebrafish left-right organ ^56–58^. *crb2a* expression was apparent in the KV precursors at tailbud stage (10 hpf) and beyond (**Figure 2G**, **Figure S2G-H**, purple arrows), and Crb2a protein (detected by indirect immunofluorescence) localized to the apical lumen of this structure in 8 somite-stage embryos (**Figure 2M-N**), consistent with localization patterns of aPKC ^64,65^, a known binding partner of Crumbs proteins ^48,66^. *crb2a* RNA was also detected in the future optic tectum (**Figure 2G, Figure S2G-H,** blue arrows), consistent with its well described roles in eye development ^35^. This anterior expression domain was unaffected in MZ*oep-/-* embryos (**Figure 1H**, **Figure S2J**) although staining was absent from the tailbud. This may indicate that Nodal does not directly regulate *crb2a* expression but, rather, is required for specification of DFCs that express *crb2a*. Alternatively, expression in this anterior domain may be regulated in a distinct Nodal-independent fashion. This analysis shows that *crb2a* is expressed in the DFCs during gastrulation, raising questions as to its role in DFCs and how these cells affect C&E morphogenesis of the adjacent notochord.

**Figure 2.**
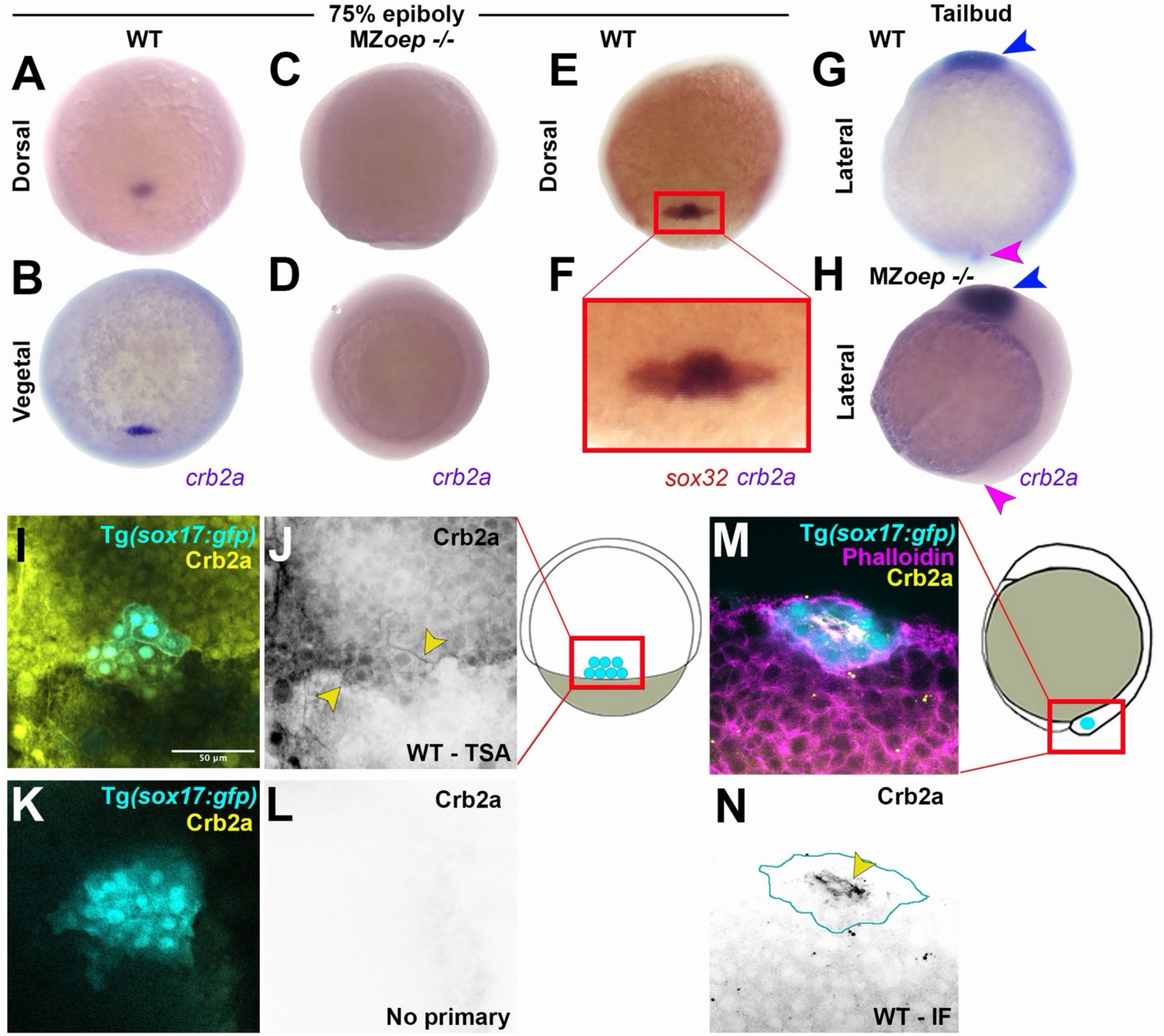
*crb2a* is expressed in dorsal forerunner cells. **A-D**) Representative images of WISH for *crb2a* at 75% epiboly (8 hpf) in WT (A-B) and MZ*oep -/-* (C-D) embryos. **E-F**) Representative images of WISH for *crb2a* (purple) and *sox32* (red) show that *crb2a-*positive cells are DFCs. **G-H**) Representative images of WISH for *crb2a* in WT (G) and MZ*oep*-/-(H) embryos at tailbud stage; *crb2a* expression in the DFCs (magenta arrowheads) but not in the optic tectum (blue arrowheads) is dependent on Nodal signaling. **I-L**) Representative images of Crb2a immunofluorescence staining with tyramide signal amplification (TSA) in WT Tg(*sox17:egfp*) embryos at 75% epiboly (I-J). Signal at the cell membranes of DFCs (J, yellow arrowheads) is absent from neighboring deep cells and from no-primary controls (K-L). J and L show the Crb2a channel from the merged images in I and K. **M-N**) Representative images of Crb2a immunofluorescence (IF) in Tg(*sox17:egfp*) sibling embryos at 8 somite-stage (13 hpf). The GFP+ Kupffer’s vesicle is outlined in Crb2a single channel image (N), yellow arrowhead highlights apical Crb2a localization in the KV. See also Figure S2 in Supplemental materials.

### *crb2a* is necessary for DFC clustering during epiboly

We next examined the effect of *crb2a* deficiency on DFCs. Two-color WISH for *crb2a* and *sox32* at 75% epiboly stage (8 hpf) failed to detect *crb2a* expression in the DFCs of *crb2a* crispants (**Figure 3A**), but also revealed a striking failure of DFCs to cluster (**Figure 3A-B**). This DFC scattering was also apparent within live *crb2a*-/-gastrulae expressing a *sox17:egfp* transgene ^67^, which labels both endoderm and DFCs (**Figure 3C-D, Video S1**). Time-lapse imaging showed that in *crb2a*-/-embryos, these cells remained scattered throughout most of gastrulation and only came together as a cluster as the yolk plug closed at gastrulation’s end (**Figure 3D)**, demonstrating that *crb2a* is required for DFC clustering during epiboly. We further observed that regions of the dorsal margin in contact with the scattered DFCs appeared to be pulled vegetally by these connections (**Figure 3D, Video S1**, yellow arrows) while neighboring DFC-free regions of the margin lagged animally. These “pulling” and “lagging” regions were reflected by reduced convexity of the embryonic margin in *crb2a*-/-gastrulae compared with sibling controls (**Figure 3E**). We also observed a widened gap between the EVL and the underlying deep cells during epiboly in *crb2a-/-* embryos compared with sibling controls (**Figure 3F-G**), suggestive of impaired mechanical coupling between these two tissue layers. Strikingly, this widened gap was specific to the dorsal side of these embryos (the normal site of DFCs), with no difference observed on their lateral sides (**Figure 3G**). This suggests that failure of DFC clustering leads to reduced coupling of the EVL and deep cells during epiboly, but only in the dorsal region.

We further observed evidence of disrupted KV function in *crb2a* deficient embryos (**Figure S3**). First, we observed smaller KVs comprised of fewer cells in *crb2a-/-* embryos at the 8-somite stage (**Figure S3A-B**), consistent with impaired DFC clustering ^68,69^. At 36 hpf, WISH for the cardiac marker *myl7* revealed a significant increase in right-ward and un-looped heart tubes upon loss of *crb2a* (**Figure S3C-D**), indicative of laterality defects that often result from KV loss or disfunction ^37,70^. Because *crb2a* also directly regulates heart morphogenesis ^37^, these heart looping phenotypes are compounded by the morphogenetic effects of impaired trabeculation, but the laterality defects remain clear (**Figure S3**). This demonstrates that Crb2a functions within DFCs to promote their clustering and, thereby, proper formation and function of the KV. We next asked whether this scattering of DFCs in *crb2a* deficient embryos is responsible for reduced C&E of the notochord, and how it may mediate this effect cell non-autonomously.

**Figure 3.**
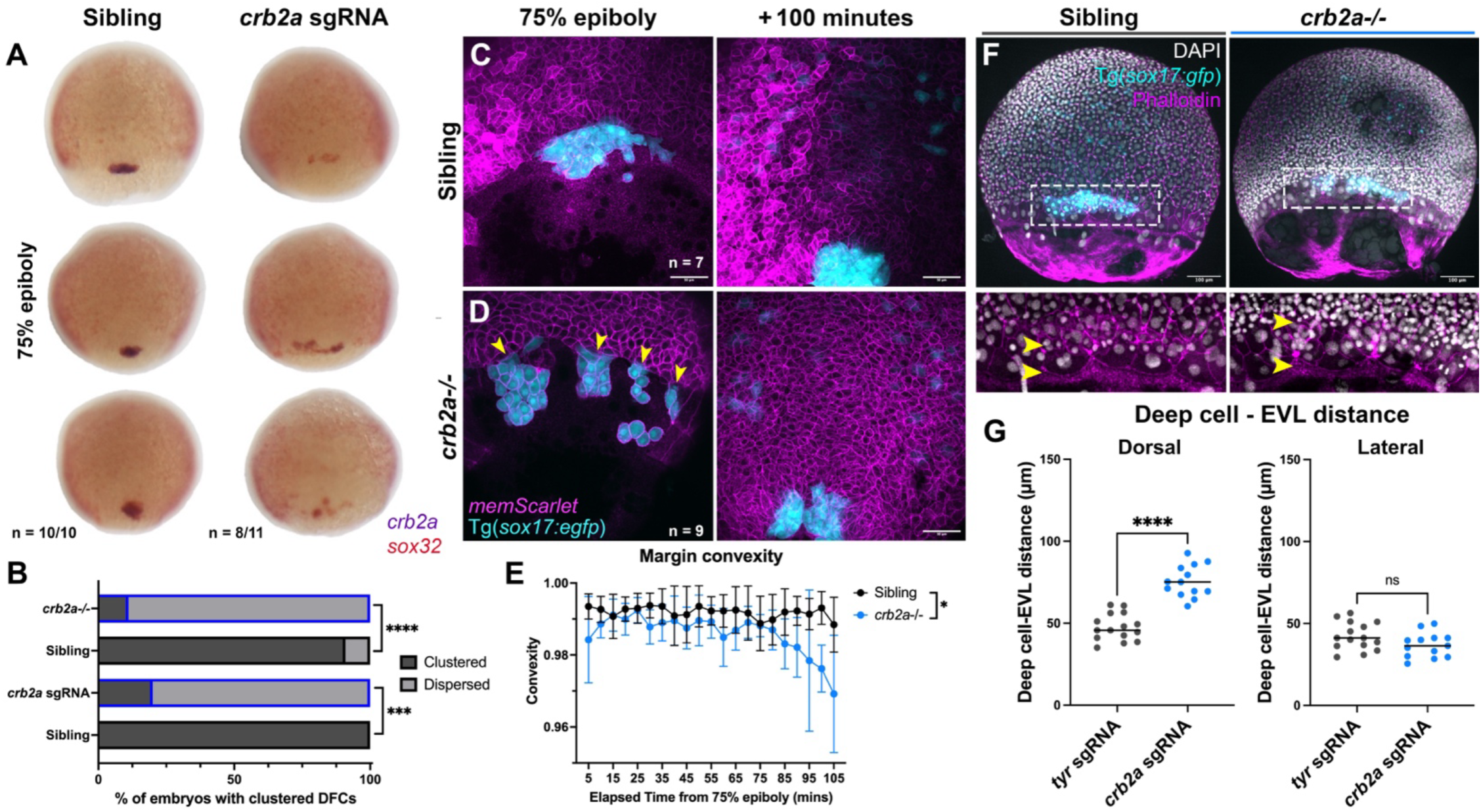
Failure of DFC clustering in *crb2a* deficient embryos causes uncoupling of deep cell and EVL epiboly. **A**) Representative images of WISH for *crb2a* (purple) and *sox32* (red) in uninjected (top) and *crb2a* crispant (bottom) embryos at 75% epiboly. Fractions indicate the number of embryos with the pictured phenotype over the number of embryos examined. **B**) Quantification of DFC clustering in *crb2a* crispants and stable mutants shown in A and C at 75% epiboly compared with sibling controls. Fisher’s exact test, *** = P < 0.001, **** = P < 0.0001. **C-D**) Representative images of live Tg(*sox17:egfp*);*crb2a*-/-mutant (D) and sibling embryos (C) at 75% epiboly (left) and 100 minutes later (right). n= number of embryos examined for each condition, arrowheads indicate regions of the margin attached to scattered DFCs. **E**) Convexity of the embryonic margin from time-lapse images of *crb2a*-/- and sibling gastrulae shown in (C-D). Symbols are means with SD. Data were analyzed by mixed effects model, P=0.0217. **F**) Representative images of DAPI and phalloidin staining in Tg(*sox17:egfp*);*crb2a* -/- and sibling embryos at 75% epiboly. Bottom images are enlarged from the regions in dashed rectangles, yellow arrowheads indicate the distance between the enveloping layer (EVL) and deep cell layers. **G**) Quantification of the distance between the EVL and deep cell layers in the dorsal (left) and lateral (right) regions of the embryos shown in (F). Each dot represents a single embryo, black bars indicate median values. T-test, **** = P < 0.0001). See also Video S1 and Figures S3 and S4 in Supplemental materials.

### DFCs promote proper epiboly and notochord C&E during gastrulation

To test whether DFCs are required for proper notochord C&E, we eliminated DFCs (and endoderm) using an antisense morpholino oligonucleotide (MO) against *sox32* ^71^ (**Figure 4A**) and performed morphometric measurements at the end of gastrulation (**Figure 4B-C**). Like *crb2a* crispants and mutants, *sox32* morphants devoid of DFCs exhibited modestly but significantly reduced AP axis length and increased notochord width without affecting the neural plate (**Figure 4C**). To rule out the possibility that these C&E defects result from loss of endoderm, we also injected *sox32* MO into the yolk cell at the 512-cell stage. Because DFCs arise from so-called Wilson’s cells, the marginal-most blastomeres that remain open to the yolk syncytial layer (YSL) through the 512-cell stage ^62^, such injections can be used to target DFCs specifically. Indeed, although DFCs were not detected in 512-cell-injected Tg(*sox17:egfp)* gastrulae, these embryos retained GFP expression in the endoderm (**Figure 4A** yellow arrowheads) and did not exhibit the characteristic cardia bifida phenotype (**Figure S3E**) seen in *sox32* morphants injected at the 1-cell stage ^71^. Like morphants injected at the 1-cell stage, these embryos showed modest but significant C&E defects (**Figure 4C**), further implicating DFCs in extension of the notochord during gastrulation. Finally, we tested whether loss of *crb2a* exacerbates C&E defects in embryos devoid of DFCs by injecting *crb2a* or control *tyrosinase* sgRNAs into *sox32* morphants. Morphometric measurements showed that axis length and notochord width in *crb2a/sox32* double deficient embryos were not significantly different than *sox32* morphants alone (**Figure 4D**). Together, these findings indicate that DFCs, and not *crb2a* per se, are essential for proper notochord C&E during gastrulation.

In addition to C&E defects, we observed increased distances between the EVL and deep cell layers of both fixed (**Figure 4E-F**) and live (**Figure S4A-B, Video S2**) *sox32* morphant embryos as they underwent epiboly. As in *crb2a*-/-embryos, this widened gap was specific to the dorsal (but not lateral) margin, highlighting a role for DFCs in coupling of dorsal tissues during epiboly. We also frequently observed blastopore closure defects in *crb2a* and *sox32* deficient embryos with scattered or absent DFCs, respectively. Although expression of a single stripe of *egr2b* in rhombomere 3 (which appears at tailbud stage) indicates that all embryos examined were at the same stage, a significantly larger number of *crb2a* and *sox32* deficient embryos possessed open blastopores (**Figure S4C-D**), indicating delayed or abnormal epiboly. Together, these observations show that epiboly movements in the dorsal deep cells lagged behind those of the overlying EVL when DFCs were dispersed or absent. We and others have found that DFCs maintain attachments to both the EVL and the posterior end of the developing notochord throughout gastrulation (**Figure S5**) ^72,73^. We therefore hypothesized that in the absence of this link, epiboly forces in the EVL are not correctly transmitted to the notochord, reducing notochord extension and epiboly progression.

**Figure 4.**
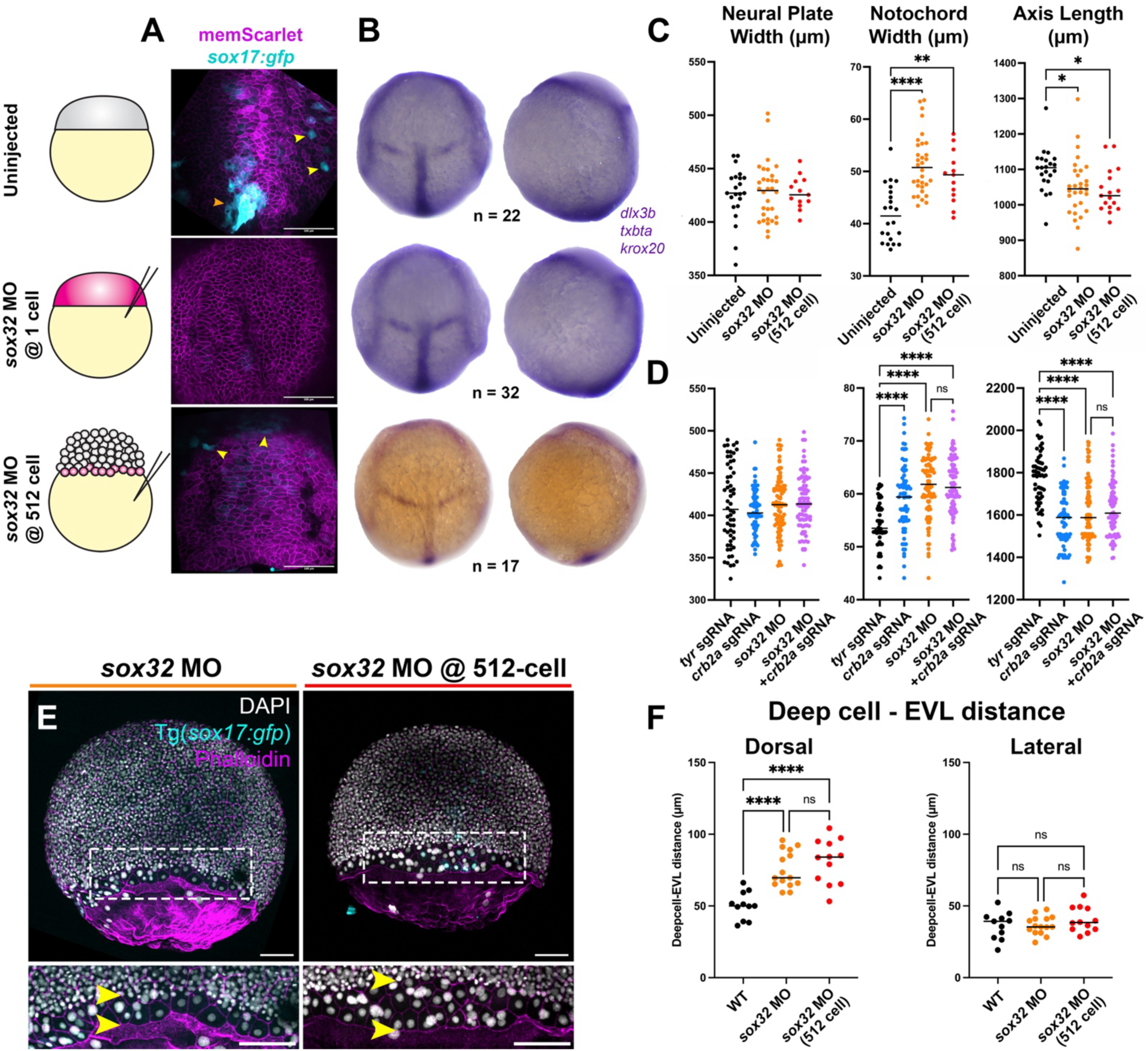
Embryos without DFCs have shortened body axes and wider notochords. **A**) Representative images of Tg(*sox17:egfp*) gastrulae of the conditions indicated. Yellow arrowheads indicate endoderm cells, orange arrowhead indicates DFCs in uninjected embryos, which are absent in *sox32* morphants. **B**) Representative dorsal (left) and lateral (right) images of WISH for *dlx3b, tbxta, and egr2b* in tailbud stage embryos of the conditions indicated. n= the number of embryos measured for each condition. **C**) Morphometric measurements of embryos shown in B. Embryos without DFCs have reduced axis length (middle) and increased notochord width (right), but neural plate is unaffected (left). Each dot represents a single embryo, black bars indicate median values. Data were analyzed by one-way ANOVA, * = P < 0.05, ** = P < 0.01, **** = P < 0.0001. **D**) Morphometric measurements as in (C) of the conditions indicated. Loss of *crb2a* did not exacerbate the reduced axis length (right) or increased notochord width (middle) phenotypes in *sox32* morphants without DFCs; neural plate remains unaffected (left). Data were analyzed by one-way ANOVA, **** = P < 0.0001. **E**) Representative images of DAPI and phalloidin staining in Tg(*sox17:egfp*) embryos injected with *sox32* morpholino at 1-cell (left) and 512-cell stage (right) at 75% epiboly. Bottom images are enlarged from the regions in dashed rectangles, yellow arrowheads indicate the distance between the EVL and deep cell layers. **F**) Quantification of the distance between the EVL and deep cell layers in the dorsal (left) and lateral (right) regions of the embryos shown in (E). Each dot represents a single embryo, black bars indicate median values. Data were analyzed by one-way ANOVA with Tukey’s post-hoc comparison, **** = P < 0.0001. See also Video S2 and Figures S3 and S4 in Supplemental materials.

### DFCs are required for polarization of posterior notochord cells

To examine whether and how disrupted DFC clustering affects cell behaviors underlying notochord C&E, we performed live time-lapse confocal imaging in Tg(*sox17:egfp*); *crb2a*-/- and sibling control gastrulae expressing membrane-localized mCherry or mScarlet. We employed Cellpose 2.0 ^74^, a machine-learning based image analysis tool, to segment individual notochord cells and FIJI ^75^ to automatically measure their elongation (aspect ratio), ML alignment (orientation of the cell’s major axis), and position of the centroid throughout gastrulation (**Figure 5A-B**). In WT and *crb2a+/-* sibling embryos, cells within the developing notochord became increasingly elongated and ML aligned throughout gastrulation, as previously reported ^9,76^. We further divided cells according to their position along the AP axis, binning them into 30 µm “margin”, margin-“adjacent”, or larger “far” regions based on their distance from the embryonic margin (**Figure 5A-B, Video S3**). Within sibling control gastrulae, cells at the embryonic margin (furthest posterior) were poorly elongated and randomly aligned throughout most of gastrulation (**Figure S6E-E’**). Those cells far from the margin (furthest anterior) were already somewhat polarized at 75% epiboly (when C&E begins ^76,77^) and became more elongated and ML aligned with time (**Figure 5C’-D’**). Cells adjacent to those in the margin, between the other two populations, were more ML aligned and elongated than margin cells but less so than “far” cells (**Figure 5C-D**). This suggests either that C&E cell behaviors within the developing notochord progress from anterior to posterior (as in *Xenopus* ^18^), that the margin and/or DFCs specifically affect the behavior of nearby cells, or both.

Within *crb2a*-/-gastrulae, we observed no significant differences in the elongation or alignment of marginal or “far” cells (**Figure S6E-E’**, **Figure 5C’-D’**), but detected striking differences in cells adjacent to the margin. Whereas sibling “adjacent” cells showed a dramatic increase in cell elongation toward the end of gastrulation, *crb2a*-/-“adjacent” cells showed essentially no increase in elongation at this time (**Figure 5C**). We also observed small but significant differences between *crb2a*-/- and sibling ML cell alignment (**Figure 5D-D’**), but it is unclear that these differences are biologically relevant. Consistent with this reduced ML cell polarization and with the results from our morphometric measurements in **Figure 1**, we found that the notochords of *crb2a*-/-embryos were significantly wider than their siblings in both the “adjacent” and “far” regions throughout gastrulation (**Figure 5E-F**).

**Figure 5.**
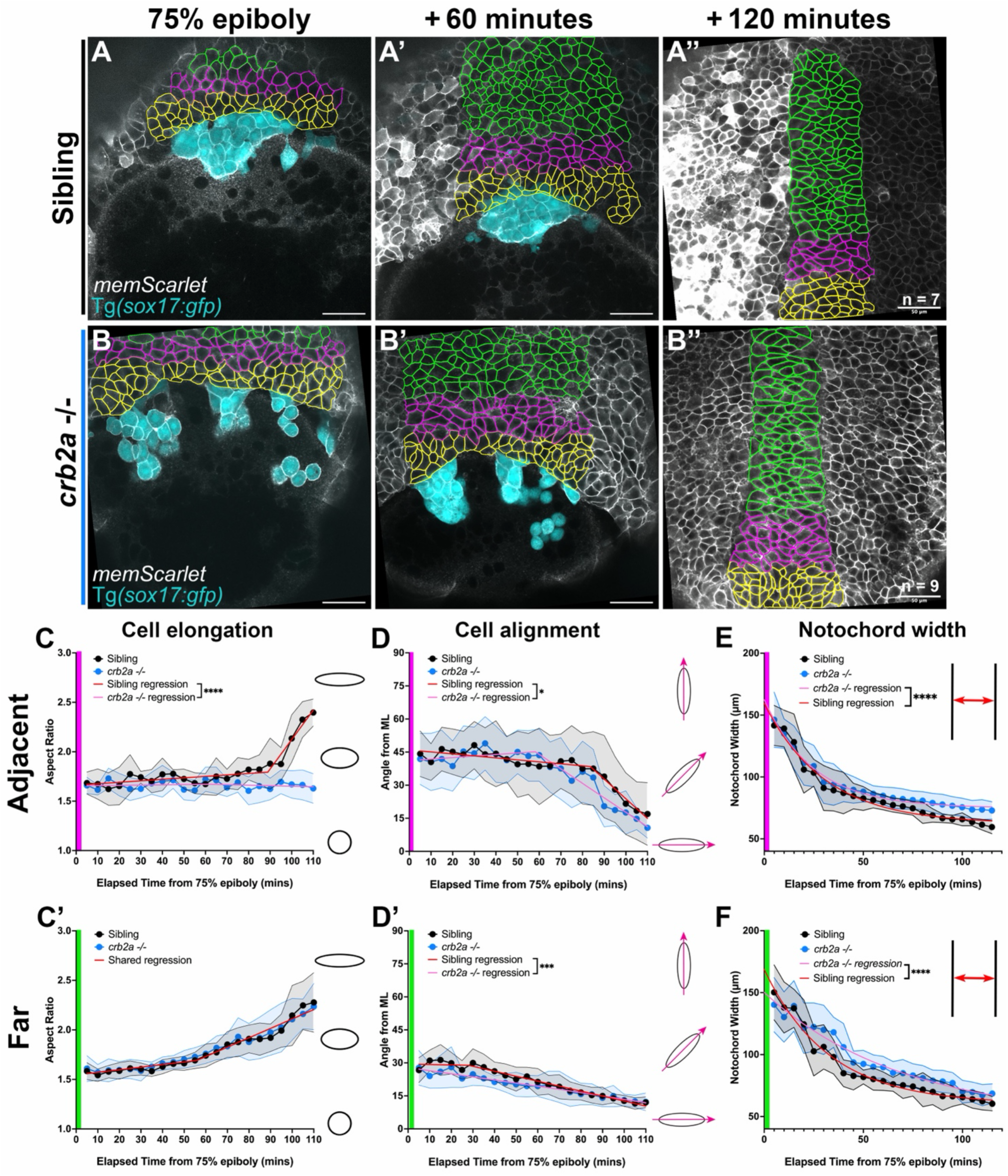
Mediolateral cell polarization is reduced in *crb2a* mutant notochords. **A-B**) Representative images from time-lapse series of the developing notochord from 75% epiboly to tailbud stage in embryos of the conditions and at the time points indicated. Colored outlines denote analyzed cells at the margin (yellow), adjacent to the margin (magenta), or far from the margin (green). n= the number of embryos analyzed per condition. **C-D**’) Quantification of sibling (black) and *crb2a*-/-(blue) notochord cell elongation (C) and cell alignment (D-D’) in the regions indicated. Graphs show medians and inter-quartile range. Data were fit with segmented lines using non-linear regression and differences detected by sum-of-squares F-tests, * = P <0.05, *** = P <0.001, **** = P <0.0001. **E-F)** Quantification of notochord width over time in the regions indicated. Data were fit with a one-phase decay non-linear regression and differences detected by sum-of-squares F-tests, **** = P <0.0001. See also Video S3 and Figures S5 and S6 in Supplemental materials.

**Figure 6.**
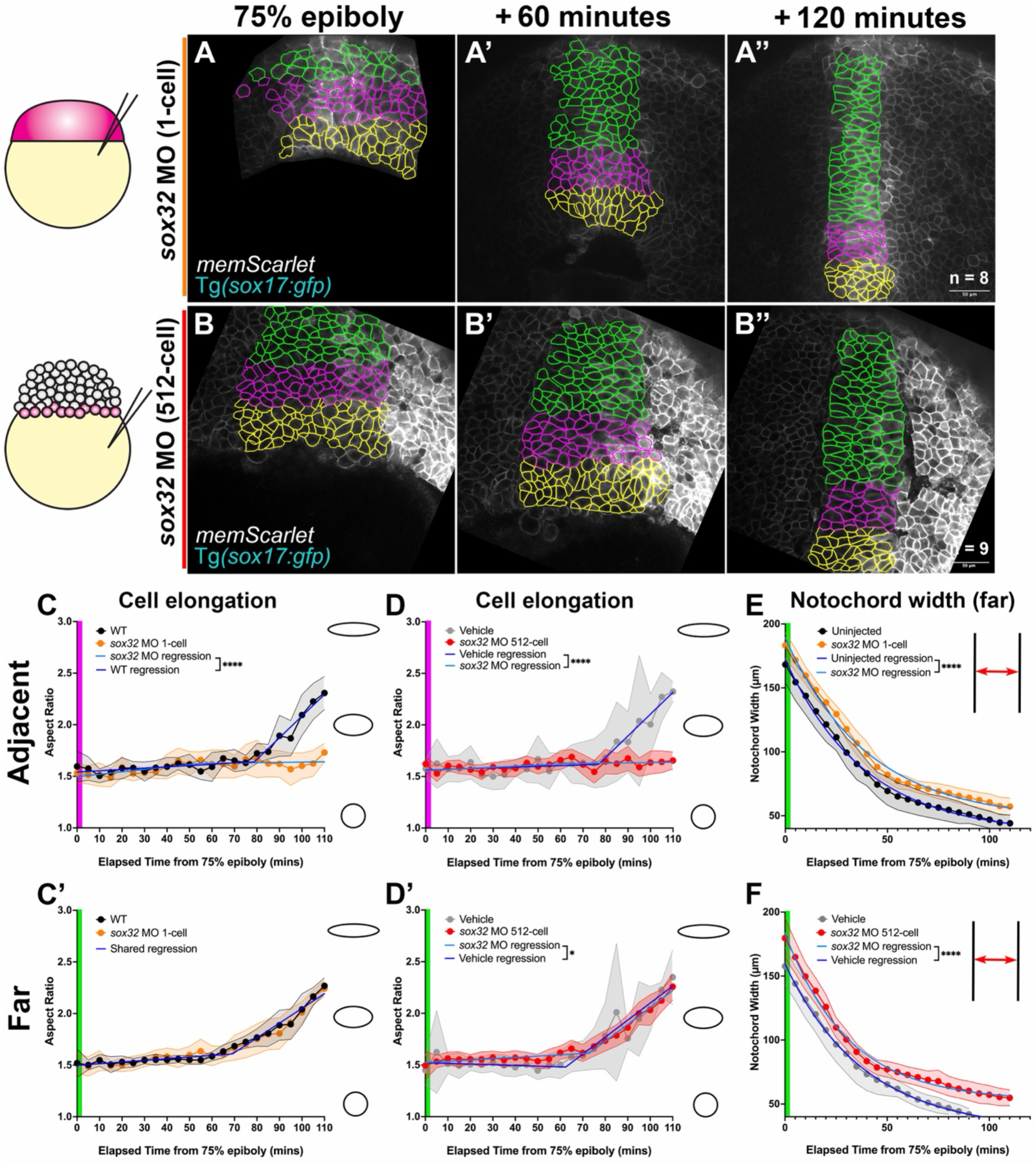
Notochord mediolateral cell polarization is reduced in *sox32* morphants without DFCs. **A-B**) Representative images from time-lapse series of the future notochord from 75% epiboly to tailbud stage in embryos of the conditions and at the time points indicated. Colored outlines denote analyzed cells at the margin (yellow), adjacent to the margin (magenta), or far from the margin (green). n= the number of embryos analyzed per condition. **C-D’**) Quantification of notochord cell elongation in the regions indicated within embryos injected with *sox32* MO at the 1-cell (C-C’, orange) or 512-cell (D-D’, red) stages and their respective controls (black and gray, N=5). Graphs show medians and inter-quartile range. Data were fit with segmented lines using non-linear regression and differences detected by sum-of-squares F-tests, * = P < 0.05, **** = P <0.0001. **E-F**) Quantification of notochord width over time in the regions indicated. Data were fit with a one-phase decay non-linear regression and differences detected by sum-of-squares F-tests, **** = P <0.0001. See also Figure S6 in Supplemental materials.

To determine if the absence of DFCs similarly affected notochord C&E cell behaviors, we performed live confocal imaging as above in embryos injected with *sox32* MO at the 1- and 512-cell stages, and in vehicle controls injected with water at the 512-cell stage (**Figure 6A-B”**). Cell shape analysis in both MO conditions revealed cell polarization defects similar to those observed in *crb2a-/-* gastrulae: cells adjacent to the margin cells failed to elongate (**Figure 6C-D**), whereas marginal and “far” cells elongated normally (**Figure 6C’-D’, Figure S6F-G).** Like our observations for *crb2a*-/-embryos, small but statistically significant changes in cell alignment resulted from *sox32* morpholino injections at the 512-cell stage (**Figure S6A-B’**), but we suspect these are not biologically relevant. Notochord narrowing was also significantly impaired in both *sox32* morphant conditions (**Figure 6 E-F, Figure S6C-D**), again consistent with our cell shape and morphometric analyses (**Figure 4C**). Taken together, these results demonstrate that clustered DFCs are necessary for the polarized cell behaviors driving C&E within the posterior notochord, likely through coupling to epiboly movements of the EVL.

### *crb2a* promotes C&E via coupling of DFCs and epiboly movements

Our findings so far demonstrate that the presence and clustering status of DFCs affects both epiboly and C&E cell behaviors within the adjacent posterior notochord, leading us to hypothesize that DFCs transmit epiboly forces from the EVL to the notochord to promote its extension. If true, we would expect that in the absence of epiboly, loss or scattering of DFCs would have no effect on the ability of tissue-intrinsic cell intercalation to drive C&E. To test this prediction, we made use of zebrafish embryonic explants which, similar to *Xenopus* animal cap explants ^27^, exhibit ex vivo C&E upon activation of Nodal signaling ^26,78^. Explants faithfully recapitulate C&E movements and cell behaviors ^26^, but because they are generated by isolating relatively naïve blastomeres from the yolk and embryonic margin ^79^, they exhibit no epiboly movements. We showed previously that these explants express *crb2a* and *sox32* ^26,34^, and by examining them for expression patterns of multiple notochord and DFC markers (**Figure S7**), we determined that they frequently exhibit clusters of DFC-like cells attached to the distal ends of notochord-like structures.

To determine the role of DFC-like cells in explant extension, we generated explants expressing the constitutively active Nodal receptor *CA-acvr1b*\* (to induce their extension), plus *sox32* MO, *crb2a* sgRNAs, or control *tyrosinase* (*tyr*) sgRNAs (**Figure 7A-E**). We fixed these explants at 12 hpf, stained them by WISH for *tbxta* to confirm the presence of notochord-like structures (**Figure 7B-E**), and quantified their length-width (L/W) ratios as a measure of C&E. We found no significant difference between L/W ratios of control *CA-acvr1b*\* *tyr* crispant, *crb2a-,* and *sox32*-deficient explants, indicating no detectable defects in C&E (**Figure 7A**). Using Tg(*flh:kaede; sox17:egfp*) double transgenic explants to simultaneously visualize notochord- and DFC-like structures in live explants (**Figure 7F**), we further showed that neither total extension (**Figure 7G**) nor notochord-specific extension (**Figure 7H**) was affected by loss of *crb2a* or *sox32*. In fact, explants with no or few DFCs extended better than those with large DFC clusters, regardless of their *crb2a* or *sox32* status (**Figure 7I**), although number of DFCs had no effect on notochord-specific extension in explants (**Figure 7J**). Together, these findings demonstrate that loss of DFCs has no effect on C&E in the absence of epiboly, supporting a model in which epiboly forces are transmitted from the EVL to the posterior end of the notochord by DFCs that physically link the two (**Figure 7K**). In the absence of this link, C&E is disrupted throughout the notochord and AP axis extension is reduced, but DFCs play no detectable role in C&E absent epiboly movements.

**Figure 7.**
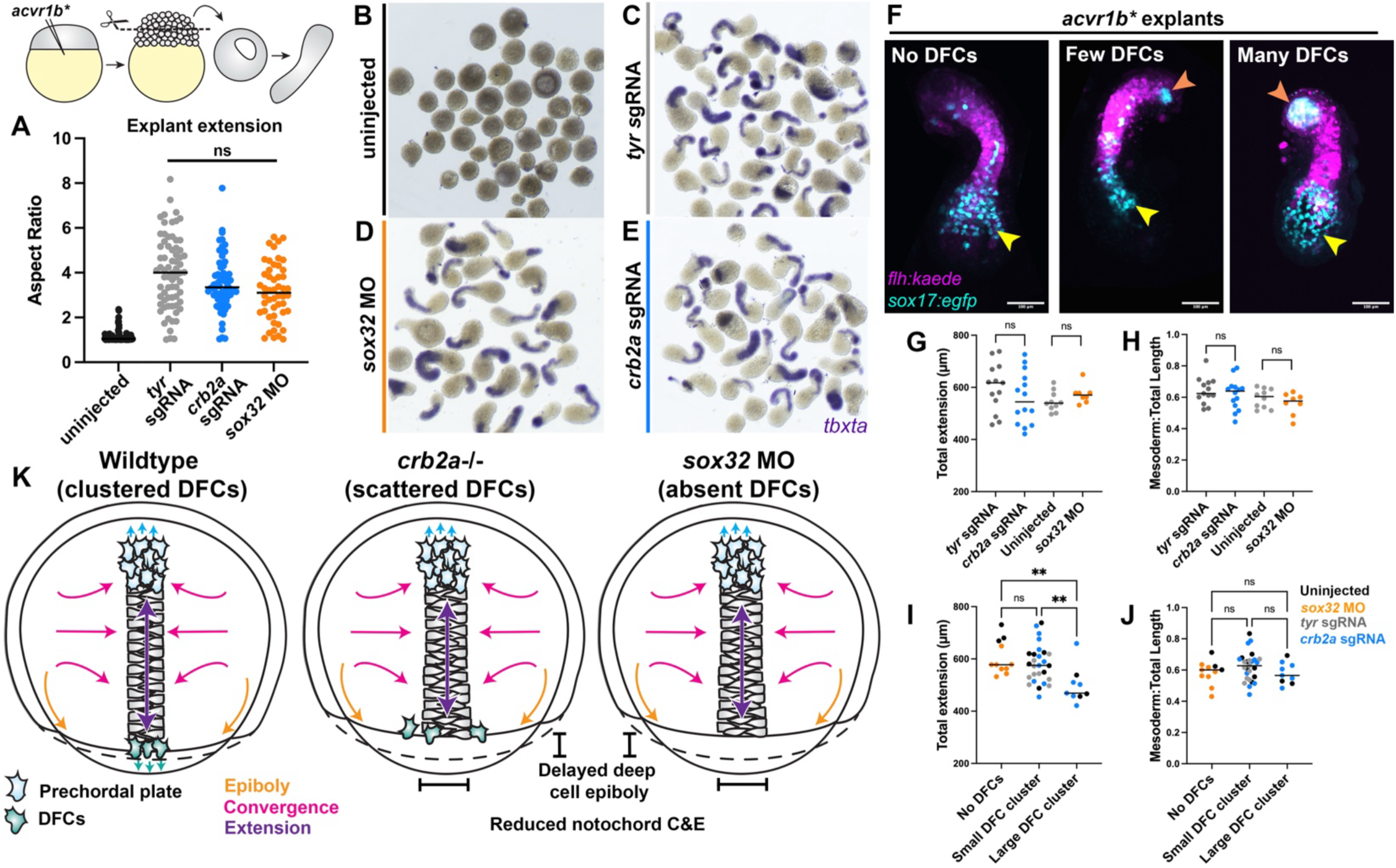
DFCs do not influence C&E in the absence of epiboly. **A**) Extension of zebrafish embryonic explants (shown in B-E) cut from *sox32* or *crb2a* deficient embryos is not affected compared with control *tyr* crispant explants. Each dot represents a single explant, bars indicate median values. Data were analyzed with the Kruskal-Wallis test, ****= P < 0.0001. Diagram is modified from ^26^. **B-E**) Representative images of *tbxta* WISH-stained embryonic explants of the conditions indicated. **F)** Representative images of double transgenic live explants expressing *flh:kaede* within notochord-like structures and *sox17:egfp* within endoderm (yellow arrows) and DFC-like cells (orange arrows). Note that DFCs frequently cluster at one pole of the explant. **G-H**) Total length (G) and ratio of mesoderm length/total length (H) in double transgenic explants lacking *crb2a* or *sox32.* Data analyzed by T-test. **I-J)** Explant length (I) and ratio of mesoderm length/total length (J) clustered by size/number of DFCs regardless of *crb2a* or *sox32* status. Data analyzed by one-way ANOVA with Tukey’s post hoc comparison, ** = P <0.01. **K)** Model for the role of DFCs in notochord C&E. DFCs enable mechanical coupling of epiboly forces to the notochord during gastrulation, enhancing its C&E. When DFCs are scattered or absent, notochord C&E and underlying cell polarization are reduced. See also Figure S7 in Supplemental materials.

## DISCUSSION

C&E morphogenesis during gastrulation is essential for proper formation of the vertebrate body plan and closure of the neural tube. Although several signaling pathways are well known to regulate the polarized cell behaviors that drive C&E, the importance of cell-extrinsic forces in this process is increasingly apparent. Here, we present evidence in support of a novel cell-extrinsic mechanism promoting C&E of the developing zebrafish notochord. We find that DFCs enhance notochord C&E, likely by transmitting vegetal-ward epiboly forces from the EVL to the posterior end of the notochord. This force supports cell-intrinsic ML cell alignment and elongation and (we speculate) directly pulls on or guides the notochord to encourage its extension. Importantly, DFCs are no longer necessary for full C&E in the absence of epiboly, demonstrating their role as transmitters of extrinsic force rather than generators of intrinsic force.

### Cell-extrinsic forces in morphogenesis

Polarized cell elongation, protrusions, and myosin contractility are thought to drive C&E by directly rearranging neighboring cells, thus propelling shape change tissue-intrinsically ^7,80^. Indeed, in *Xenopus* embryos from which the blastocoel roof was removed, the mesoderm extends autonomously in the absence of a migratory substrate ^81,82^. However, cell-extrinsic forces (i.e. those generated by neighboring tissues) are also well documented to contribute to axial extension. For example, mesoderm internalization in Drosophila gastrulae generates mechanical forces that promote extension of the ectodermal germ band ^13^. The closest parallel to the model described in this study, however, is the anterior migration of prechordal plate and head mesoderm of zebrafish and *Xenopus* gastrulae, respectively, that exert forces on the anterior end of the developing notochord ^14–17^. In these tissues, anterior axial mesoderm cells form a shingle-like arrangement in which the leading edge of one row of cells overlaps with the trailing edge of the cells ahead of them ^83^. The adhesion of leading cells to the trailing cells behind them is essential for their coordinated anterior migration ^84–87^, and pulling on cadherins in one direction is sufficient to reorient a cell’s migration in the opposite direction ^84^. Cell autonomy transplant and laser ablation experiments in zebrafish gastrulae demonstrated that failure of proper prechordal plate migration or loss of its connection to the notochord disrupts C&E of the trailing notochord non-autonomously ^15,85^. Combined with our findings, this supports a model in which pulling from both the anterior and posterior ends of the developing notochord is required for its full extension.

One important distinction between the anterior axial mesoderm and the DFCs as sources of these pulling forces, however, is that the prechordal plate actively migrates, while displacement of DFCs is driven by epiboly movements of the adjacent EVL ^55^. Epiboly is a well-studied morphogenetic process that also results from a combination of intrinsic and extrinsic forces. The initial stages of epiboly are driven by a fluidization of the animal blastoderm and radial cell intercalation that thins the blastoderm and spreads it over the yolk ^88,89^. Once the blastoderm reaches the approximate equator of the embryo (50% epiboly), an actomyosin cable within the yolk syncytial layer (YSL) at the embryonic margin contracts circumferentially, pulling itself and the EVL toward the vegetal pole ^90,91^. Vegetal movement of the YSL actomyosin cable can proceed when all cells have been removed from the yolk, and even moves faster than in intact embryos ^92^, suggesting that embryonic cells exert drag on this structure. Adhesion to the EVL is required for the deep cell layer to spread during epiboly, as evidenced by the increased distance between these cell layers upon loss of E-cadherin and Epithelial cell adhesion molecule (Epcam) ^93,94^.

In our model, we envision DFCs as a trailer hitch that enables these strong epiboly forces (the truck) to be transferred to the posterior notochord (the trailer), stretching the tissue anchored at the opposite (anterior) end. This is supported by the observation that the distance between the EVL and deep cells increases specifically in the dorsal region (**Figure 3F-G**, **Figure 4E-F, Figure S4A-B**) when DFCs are scattered or absent. Although notochord cells also generate intrinsic forces through cell intercalation, these additional extrinsic influences may ensure robustness of the C&E process. Indeed, we found that although loss or scattering of DFCs only disrupts cell-intrinsic ML alignment and elongation of notochord cells near the dorsal margin (**Figs. 5 & 6**), notochord convergence is reduced along its entire length. This is consistent with the recent finding that localized forces are propagated across embryonic tissues to promote C&E ^95^. Our data suggest that this “trailer hitch” mechanical cue also prevents yolk plug closure defects, ensures correct positioning of the KV, and synchronizes epiboly between cell layers, thereby increasing the likelihood of successful embryogenesis. Although we cannot rule out an alternative mechanism by which DFCs promote C&E of the adjacent notochord, our finding that DFCs have no impact on C&E in the absence of epiboly forces (**Figure 7**) strongly supports this trailer hitch model.

### Crumbs2a in gastrulation

We initially selected *crb2a* as a candidate morphogenetic effector of Nodal signaling because its loss causes C&E defects in mouse gastrulae ^49^ and its subcellular localization promotes cell and tissue anisotropy of Myosin in *Drosophila* ^47,48^. To our surprise, however, the role of *crb2a* in zebrafish gastrulation is entirely different, apparently serving only to facilitate DFC clustering. Notably, DFCs are still physically able to cluster and form a KV when brought into proximity after blastopore closure in *crb2a*-/-embryos, so this effect does not appear to be solely related to their ability to adhere to one another. Rather, we speculate that disrupting the EVL-DFC-deep cell nexus may affect the ability of the DFCs to cluster together at the midline earlier in gastrulation. It was shown that DFCs maintain connections with the EVL (from which they partially delaminate) via their narrowed apical ends ^55^, hence, an apical determinant in this process is a logical requirement. However, our data suggest that the role for Crb2a in DFCs goes beyond maintaining apical surfaces, and may involve homophilic adhesion between Crb2a extracellular domains on adjacent cells ^47,96,97^, intracellular interactions with FERM domain proteins to organize actomyosin ^47,98,99^, and/or additional functions. Although it is beyond the scope of this study, future work may determine which Crb2a domains and interacting partners are necessary for DFC clustering and, consequentially, C&E.

### A role for DFCs in gastrulation morphogenesis

Multiple experimental manipulations were found to cause DFC scattering in zebrafish gastrulae, but to our knowledge, this was not identified as the source of any additional morphogenetic defects during gastrulation. For example, cold temperatures ^68,100^, loss of Cadherins ^101^ or other cell adhesion molecules ^70^, loss of Integrins ^69^, Eph-ephrin signaling ^64^, and even PCP signaling ^72^ reduced the number and clustering of DFCs, yielding small and/or fragmented KVs and ultimately left-right patterning defects. Aside from PCP mutants, reduced C&E was not reported in these gastrulae, nor in *sox32/casanova* mutants entirely lacking DFCs ^71^. However, given that the axis extension defects in *sox32* and *crb2a* deficient embryos are relatively mild, they were likely overlooked in studies not explicitly examining C&E. It is also possible that epiboly defects in embryos with abnormal DFCs masked their effect on C&E, as embryos with open yolk plugs may have been dismissed as generally delayed. We note that most *crb2a* and *sox32* deficient gastrulae did successfully close their yolk plugs, suggesting that force transmission from the EVL to the deep cells is not strictly necessary to complete epiboly. Instead, we suspect that that this mechanical cue acts as a buffering system to increase the likelihood that epiboly will complete successfully. Our work thus highlights how cell-intrinsic and -extrinsic forces cooperate to ensure developmental robustness during gastrulation.

## RESOURCE AVAILABILITY

### Lead contact

- Requests for further information and resources should be directed to and will be fulfilled by the lead contact, Margot Williams.

### Materials availability

- The *crb2a^bcm122^* zebrafish line generated in this study will be made available from the lead contact upon request.

### Data and code availability

- This study analyzes existing publicly available RNA-sequencing data accessible in GEO under accession codes GSE147302 and GSE246158.
- All original data reported in this paper will be shared by the lead contact upon request.
- This paper does not report original code.
- Any additional information required to reanalyze the data reported in this paper is available from the lead contact upon request.

## ACKNOWLEDGEMENTS

We are grateful to Dr. Lila Solnica-Krezel for sharing plasmids and WISH probes, Dr. Lance Davidson for sharing plasmids, Drs. Rosa Uribe and Patrick Blader for sharing transgenic fish lines, and Dr. Irina Larina for 3D printing molds for mounting embryos. We also thank the BCM Center for Comparative Medicine for taking excellent care of our fish, Jacalyn MacGowan for help with immunostaining, and all members of the Williams lab for their help and feedback on this project. This work was supported by NIH/NICHD grants R00HD091386 and R01HD104784 to M.K.W.

## AUTHOR CONTRIBUTIONS

C.S. and M.K.W. conceived of the project. C.S. and M.K.W. performed experiments and analysis. C.S. and M.K.W. wrote and edited the manuscript.

## DECLARATION OF INTERESTS

The authors declare no competing interests.

## METHODS

### 1. Experimental model details

#### Zebrafish

Adult zebrafish were reared according to established protocols ^102^ in accordance with the standards set by Baylor College of Medicine Institutional Animal Care and Use Committee. Embryos were collected by natural spawning and staged according to ^103^. Experiments were conducted using embryos from AB WT, Tg(*sox17*:e*gfp*)*^s870tg^* ^67^, Tg(*flh:kaede*)*^vu376tg^* ^104^, Tg(*lhx1a:gfp*)*^pt303tg^* ^105^, and *crb2a^bcm122^* (this study, described below) strains. Embryos of the strains and stages reported within the text and figure legends were selected randomly for inclusion experiments. Because sex is not determined in zebrafish until approximately two weeks of age, sex is not a relevant biological variable in the <3 day post fertilization embryos used in this study.

### 2. Methods details

#### Generation of *crb2a* mutant zebrafish

An sgRNA targeting the last exon of *crb2a* (exon 13) was transcribed in vitro using T7 RNA polymerase (NEB M0251), pre-complexed with EnGen Spy Cas9 NLS (NEB M0646M), and injected into Tg(*sox17:egfp*) embryos at the 1-cell stage (as described below). Embryos were raised until 72 hpf when the *crb2a*-/-phenotype could be observed unequivocally to confirm the effectiveness of the sgRNA. The amount of sgRNA-Cas9 complex injected was then titrated to yield ∼50% of embryos with the expected phenotype, and the remaining non-phenotypic siblings were raised as founders. Adult F0s were in-crossed to identify pairs that produced phenotypic *crb2a*-/-embryos, which were then outcrossed to WT adults to establish stable lines. The CRISPR target site was PCR amplified from F1 fin clip DNA and lesions were identified by Sanger sequencing.

#### Blastoderm explants

Blastoderm explants were generated according to ^79^. Embryos were dechorionated with Pronase (1 ml of 20 mg/ml stock in 14 ml 3x Danieau’s solution) at the 128-cell stage. After sequential washes with 0.3x Danieau’s solution and egg water, explants were cut using Dumont #55 forceps (Fisher Scientific, NC9791564) in a 60 mm x 15 mm plate coated with agarose and filled with 3x Danieau’s solution. Explants were allowed to heal in 3x Danieau’s solution for 5 minutes before transfer into agarose-coated 6-well plates filled with explant media (DMEM/F12 (Thermo Fisher Scientific, 11330032), 3% newborn calf serum, 1:200 penicillin-streptomycin).

Explants were collected and fixed in 4% paraformaldehyde (PFA) when stage-matched embryos reached the 4-somite stage. Explant extension was measured as the length/width ratio of explants after fixation, calculated by measuring the length of the major axis using the segmented line tool in FIJI ^75^ and dividing by the width at the midpoint of the explant.

#### Microinjections

Embryos were loaded into troughs of agarose molds (Adaptive Science Tools, I-34) and injected with 0.5-2 nL volumes using pulled glass needles (Fisher Scientific, 50-821-984). mRNA concentrations were measured using a Nanodrop 2000.

##### mRNAs

mRNAs were synthesized using the SP6 or T7 mMessage Machine Kits (Fisher Scientific, AM1340 or AM1344 respectively) and purified using either BioRad Microbiospin columns (Bio-Rad, 7326250) or LiCl precipitation as described in the mMessage Machine Kit protocols. For explant experiments, embryos were injected with 0.5 pg CA-*acvr1b\** ^106^. For live imaging experiments, embryos were injected with 200 pg *membrane-cherry* (gift from Dr Fang Lin, Department of Anatomy and Cell Biology at University of Iowa Carver College of Medicine, IA, USA) or 100 pg of *membrane-scarlet* ^107^ (gift from Dr Lance Davidson, Department of Bioengineering, Developmental Biology and Systems Biology at University of Pittsburgh, Pittsburgh, PA, USA).

##### sgRNAs

DNA templates for sgRNA production were generated by overhang PCR (see key resources table for all primer sequences). Target sites for Cas9 were identified using CRISPRscan ^108^ and primers for target site amplification were designed using CHOPCHOP ^109^. *tyrosinase* sgRNA sequences are from ^110^. Any further *in silico* work (ie: aligning sequencing files) was performed using ApE ^111^. sgRNAs were synthesized using T7 RNA polymerase (NEB, M0251) and purified by ammonium acetate precipitation as previously described ^59,112,113^. For F0 multi-guide CRISPR targeting, three sgRNAs per gene were synthesized, complexed with EnGen SpyCas9 (NEB, M0646) *in vitro,* and injected (1 nl volume) at the 1-cell stage as described in ^59^ for each of the following genes: *crb2a, vangl2,* and *tyrosinase*.

##### Morpholino oligonucleotides

A well-characterized morpholino against *sox32* (MO1-*sox32* ^114^, Gene tools) was injected either at the 1-cell stage (targeting the embryo *in toto*) or at the 512-cell stage (targeting Wilson’s cells) into Tg(*sox17*:*egfp*) embryos as described in ^115^. Injection success was assayed by observing injected embryos on a Nikon SMZ18 epifluorescence microscope between 50% and 60% epiboly; embryos successfully injected at the 1-cell stage had no detectable GFP in either endoderm or in the DFCs, whereas embryos successfully injected at the 512-cell stage displayed GFP in the endoderm but no patch of GFP at the margin.

##### Immunofluorescence staining

Embryos and explants were fixed in 4% PFA at the desired stages, washed with PBSTr (PBS with 0.1% Triton X100), blocked overnight at 4°C (PBSTr with 10% newborn calf serum), incubated overnight with a primary antibody raised against zebrafish Crb2a ^96^ (undiluted) or Laminin (Millipore-Sigma L9393, 1:500), washed in PBSTr, and then incubated overnight at 4°C in a 1:1000 secondary antibody (Invitrogen, A32728) solution containing 1:10000 DAPI (ThermoFisher Scientific, 62248) and 1:1000 rhodamine phalloidin (ThermoFisher Scientific, A22283). For embryos at 75% epiboly, a cyanine 3 tyramide signal amplification (TSA) kit (Akoya biosciences NEL744001KT) was used to amplify Crb2a staining. Embryos were fixed in 4% PFA at the desired stage, washed with PBST (PBS with 0.1% Tween), blocked for 1 hour at RT with 10% fetal bovine serum in PBT, and incubated overnight at 4°C with undiluted Crb2a antibody. Embryos were then washed extensively in PBST and incubated overnight at 4°C with an anti-Mouse HRP antibody (Bio-Rad #1706516) diluted 1:500 in block. After washing extensively in PBST, embryos were incubated for 15 minutes in TSA amplification buffer, then in tyramide working solution for 45 minutes. Staining was stopped by incubating with Reaction Stop Reagent for 1 hour and washed with PBST before imaging.

#### Whole mount *in situ* hybridization

Antisense riboprobes were synthesized using NEB T7 RNA polymerase (NEB, M0251) and digoxigenin NTPs (Millipore Sigma, 11277073910) or fluorescein NTPs (Millipore Sigma, 11685619910). *In situ* hybridization was performed according to Thisse and Thisse ^116^ with the following modifications for fluorescein probes: embryos were washed in MABT (100 mM maleic acid, 150 mM NaCl, 0.1% Tween-20, pH 7.5) and incubated in RBR buffer (20% goat serum, 2% Roche blocking reagent (Millipore Sigma, 11096176001) in MABT) overnight at 4°C on a shaker. Embryos were then washed in MABT before 3 5-minute incubations in staining buffer (100mM Tris (pH 9.5), 50mM MgCl2, 100 mM NaCl in PBT (PBS with 0.1% Tween-20)). Staining buffer was removed and embryos were incubated in staining solution until the stain was detectable, at which time embryos were washed into 10 mM EDTA in PBT to stop the staining reaction. WISH stained embryos were mounted in 3% methylcellulose for imaging.

#### Genotyping

Adult fin clips and gastrulae from *crb2a+/-* in-crosses were genotyped by PCR using allele-specific primer pairs (see key resources table) that amplify from only WT or *bcm122* exon 13. DNA from WISH-stained embryos was extracted using a high-salt de-crosslinking step prior to genotyping as described previously ^117^. Briefly, embryos were incubated at 65°C for 4 hours in 300 mM NaCl to reverse crosslinking. NaCl solution was removed and replaced with 50 µl embryo lysis buffer (500 µl 1M Tris pH 8.0, 2.5 ml 1M KCl, 150 µl Tween-20, 47 ml water) and 4 µl Proteinase K. Embryos were incubated for 8 hours at 55°C and 10 minutes at 98°C to inactivate the proteinase K before storage at -20°C until genotyping. Embryos subjected to live confocal imaging were isolated after completion of imaging and raised to 48 hpf to identify those with *crb2a-/-* phenotypes.

#### Microscopy

Confocal microscopy was performed using a Nikon ECLIPSE Ti2 confocal microscope fitted with a Yokogawa W1 spinning disk unit, PFS4 camera, 405/488/561/633 nm lasers and 10x, 20x, and 40x Plan Apo Lambda lenses. For live imaging experiments, embryos were maintained at 28°C using a Tokai Hit STX stage heater. Embryos were mounted in glass-bottomed Petri dishes (Fisher Scientific, FB0875711YZ) coated in 0.3% (in egg water) low melting point agarose (Thermo Fisher Scientific, 16520100). Time-lapse series for cell shape analysis were collected using the 40x lens with a 2-µm step size and a 5-minute time interval for 3 hours. For experiments that required post-hoc genotyping (ie: *crb2a*-/-stable mutants), embryos were excised from the agarose and identified by phenotype. Tg(*flh:kaede*) embryos and explants were photo-converted with a 10-second pulse of 405 nm illumination before imaging. Fixed embryos were mounted in custom 3D printed agarose (1.2% agarose in egg water) mold designed by Zaucker *et al.* ^118^. Molds were generously printed by the lab of Dr. Irina Larina (Department of Integrated Physiology, Baylor College of Medicine, Houston, TX, USA) on a liquid 3D printer (Form2, Formlabs, Somerville, MA, USA). Bright-field images of live and fixed WISH-stained embryos were collected using a Nikon Fi3 color camera on Nikon SMZ745T stereoscope.

### 3. Quantification and statistical analysis

#### Image analysis

All microscopy data sets were prepared, processed, and analyzed using ImageJ/FIJI ^75^.

##### Whole embryo morphometrics

Neural plate measurements were obtained by drawing a line between the *dlx3b* WISH domains at the position of rhombomere 3 marked by the *egr2b* domain. Notochord measurements were obtained by measuring the width of the notochord also at the *egr2b* domain. Axis length measurements were obtained by measurement the length of the *dlx3b* domain from lateral-view images. All embryo images were blinded using the Perl script *blind_renamer* (https://github.com/jimsalterjrs/blindanalysis) (blindanalysis: v.1.0.) prior to analysis to ensure unbiased measurements. Embryos from *crb2a+/-* in-crosses were genotyped by PCR only after measurements were taken from WISH images.

##### Cell Shape analysis

Time-lapse Z-stacks were manually curated to create single plane videos of notochord cells. These single plane videos were manually cropped to include only future notochord cells prior to segmentation to facilitate downstream automated cell shape analysis. Cropped videos were saved as images sequences (one image per timepoint) and fed into CellPose 2.0 ^74^ with Python for segmentation. Using a FIJI macro, each set of ROIs was overlaid onto its corresponding image, all ROIs were measured (including major axis length, minor axis length, angle of the major axis, total area, and Y position), and exported as a single .csv file per time point for every embryo. Data from these csv files were collated with Python, filtered by total area (>50 µm and <300 µm), sorted based on their position in Y relative to the Y position of the margin of epiboly, and imported into GraphPad Prism 10 for statistical analysis. Notochord widths were measured in FIJI by working backwards through the videos from time points when the notochord boundaries could be clearly observed and tracking those chordamesoderm cells to their initial positions at the start of the videos.

##### Margin convexity analysis

Z-projections were made time-lapse confocal image series of membrane labeled embryos from Tg(*sox17:egfp*); *crb2a*+/-incrosses to include as much of the embryonic margin as possible. Researchers were blinded to embryo genotype as above. FIJI was used to draw a polygon selection around the margin of the deep cells, excluding DFCs, at each time point and its perimeter measured. A convex hull was then fit to each polygon, and its perimeter was measured. The convex hull perimeter was then divided by the margin perimeter to yield a convexity index for each time point.

##### Kupffer’s vesicle analysis

*crb2a* heterozygous adults (in a Tg(*sox17:egfp*) background) were incrossed and embryos were fixed at the 8-somite stage overnight at 4°C in 4% PFA. Embryos were washed with 0.1% PBSTr, stained overnight at 4°C with DAPI (1:10000) and rhodamine-phalloidin (1:1000), washed again with 0.1% PBSTr, and imaged with confocal microscopy while blinded to genotype. Nuclei within the KV were counted using the MiToBo Cell Counter ^119^ and embryos were then genotyped by PCR as described above.

#### Statistical Analysis

GraphPad Prism 10 was used for all statistical analyses and to generate graphs for all data analyzed. Statistical tests were chosen appropriately based on the type and normality of the data. The specific statistical tests used for each experiment are described in the figure legends.

## SUPPLEMENTAL VIDEO TITLES AND LEGENDS

**Video S1. Convergence & extension of the notochord in embryos with clustered and scattered DFCs.** Time-lapse confocal series of Tg(*sox17:egfp)* sibling (left) and *crb2a -/-* (right) embryos from 75% epiboly to tailbud stage. Magenta channel is memScarlet, *sox17:egfp* in cyan labels DFCs and endoderm cells. Yellow arrows highlight regions of the dorsal margin in this *crb2a*-/-embryo where the adjacent DFCs appear to pull the margin vegetally compared with DFC-regions. White arrows highlight the boundaries of the nascent notochord.

**Video S2. Convergence & extension of the notochord in *sox32* morphant embryos without DFCs.** Time-lapse confocal series of Tg(*sox17:egfp)* embryos injected with *sox32* morpholino at the 1-cell stage (left) or 512-cell stage (right), from 75% epiboly to tailbud stage. Magenta channel is memScarlet, *sox17:egfp* in cyan labels DFCs and endoderm cells. White arrows highlight the boundaries of the nascent notochord.

**Video S3. Image analysis pipeline – cell segmentation and compartmentalization.** Time-lapse confocal series of a WT/sibling zebrafish embryo labeled with memScarlet from 75% epiboly to tailbud stage. Yellow, magenta, and green cell outlines denote “margin”, “adjacent”, and “far” notochord cells, respectively.

## Notes

### Competing Interest Statement

The authors have declared no competing interest.

### Summary of Updates

Additional and updated analysis is now included in Figures 2 and 3. Language was tempered regarding mechanical forces.

