## Supplemental materials for "Dorsal forerunner cells couple epiboly movements to zebrafish notochord extension"

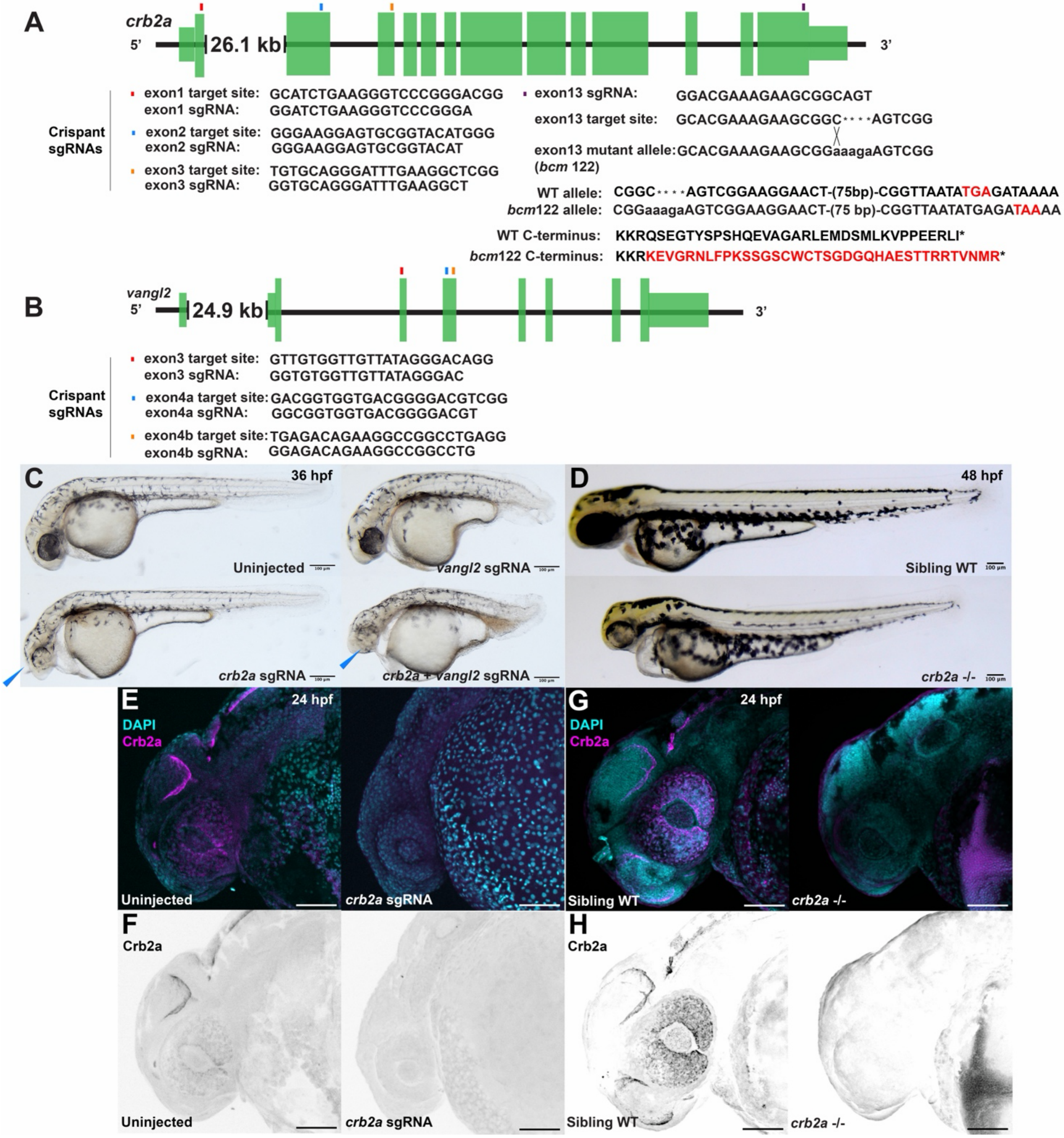


**Figure S1**. ***crb2a* crispants recapitulate the phenotype of *oko meduzy* mutants**. Related to Figure 1. **A**) Schematic of *crb2a* genomic sequence with green rectangles representing exons. Colored marks above the model denote sgRNA target sites for both crispants (exons 1, 2 and 3) and for stable *crb2a* mutant line (exon13). Target site sequences and sgRNA sequences are listed below. **B**) Schematic of *vangl2* genomic sequence, colored marks above the model denote sgRNA target sites for crispants (exons 2, 3a and 3b). Target site sequences and sgRNA sequences are listed below. **C**) Representative bright field images of uninjected embryos (top left) and crispants for *vangl2* (top right), *crb2a* (bottom left), and *crb2a* + *vangl2* double crispants (bottom right) at 36 hpf. Blue arrowheads denote the loss of eye pigmentation characteristic of *oko meduzy* mutants. **D**) Representative bright field images of sibling embryos (top) and *crb2a*-/- embryos (bottom) at 48 hpf. **E-H**) Representative immunofluorescence images of anti-Crb2a staining at 24 hpf in *crb2a* crispants and uninjected controls (E-F) and *crb2a*-/- embryos and siblings (G-H). Bottom panels are single channel images of Crb2a (F, H) from the composites in E and G.

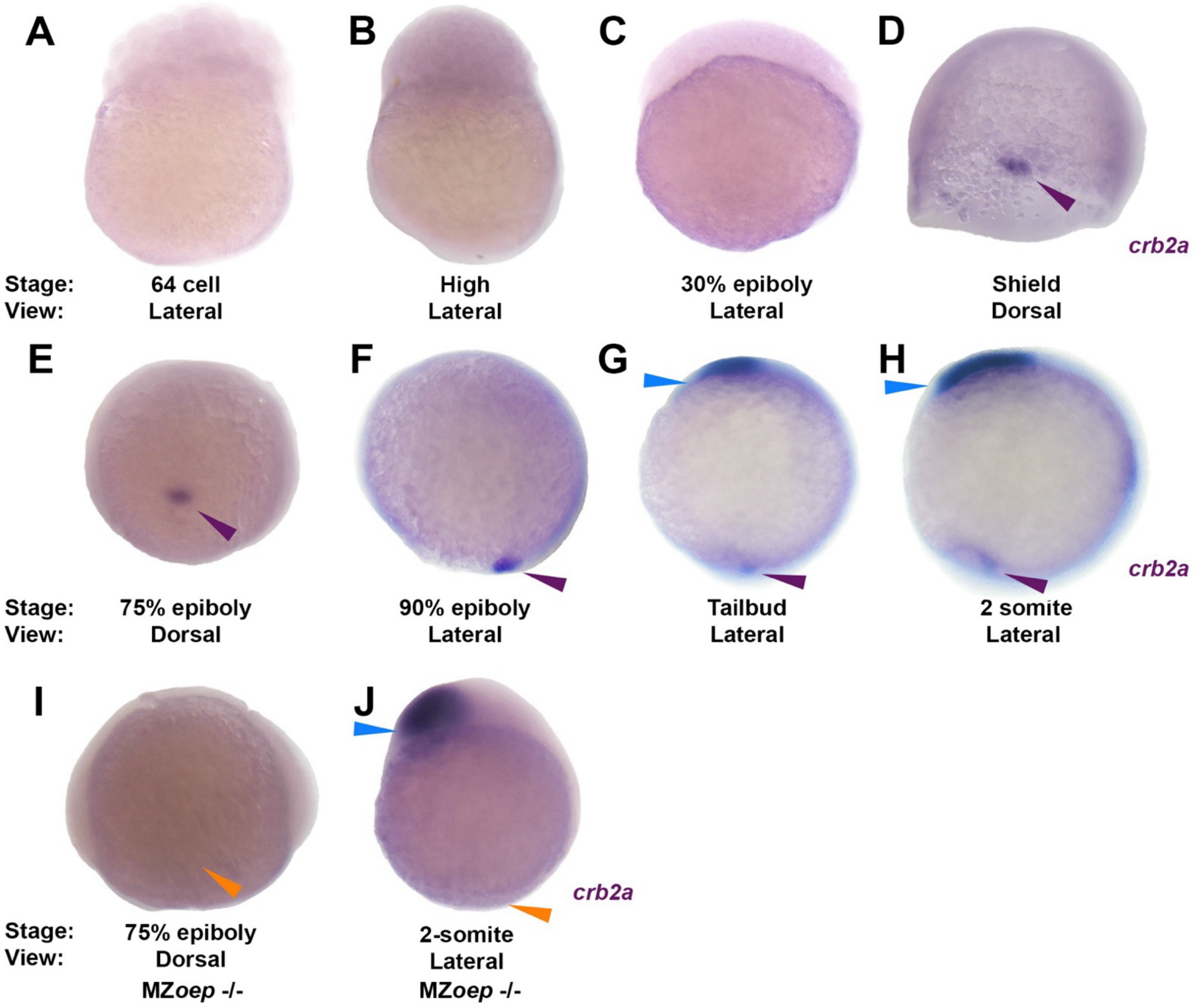


**Figure S2**. ***crb2a* expression patterns during gastrulation**. Related to Figure 2. WISH for *crb2a* in staged WT (A-H) and MZ*oep*-/- (**I**-J) embryos. **A-C**) *crb2a* is not detectable before gastrulation. **D-F**) *crb2a* (purple arrowheads) is first detected at shield stage in the dorsal forerunner cells at the dorsal midline of the margin. **G-H**) From tailbud stage onward, *crb2a* expression is also detected in the optic tectum (blue arrowheads). **I-J**) *crb2a* is not detected at the margin in MZ*oep*-/- embryos (orange arrowheads), but is expressed normally in the optic tectum (blue arrowheads).

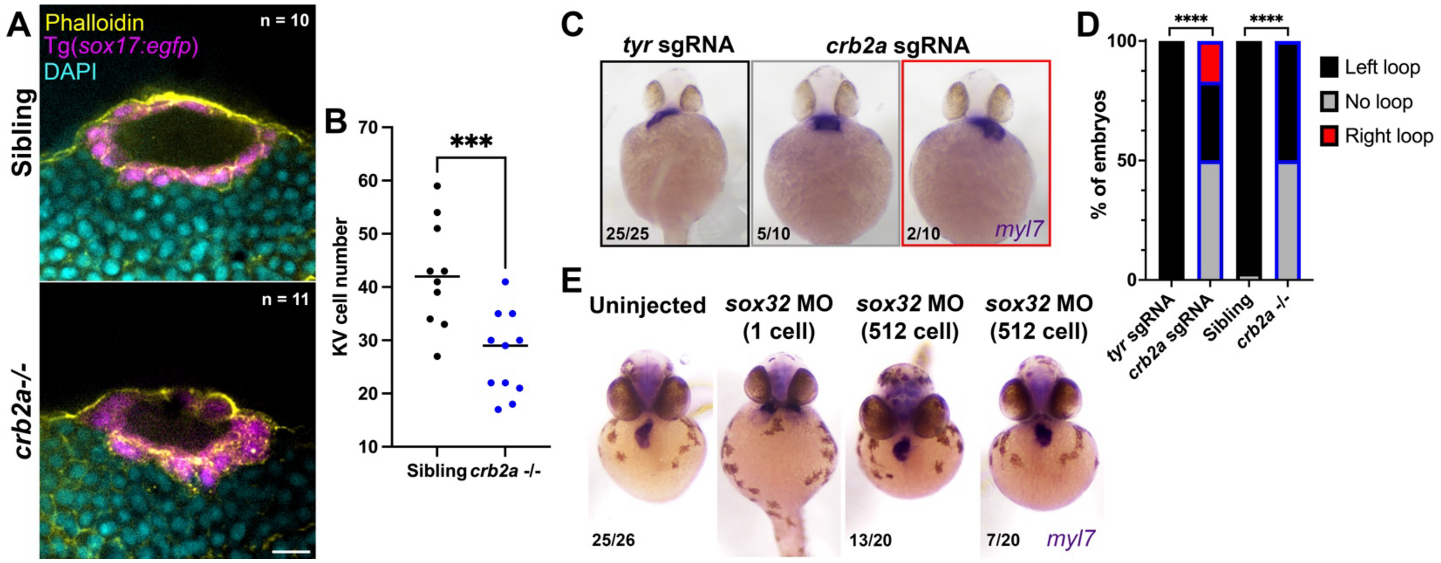


**Figure S3. Laterality defects in embryos with disrupted DFCs**. Related to Figures 3 and 4. **A**) Representative images of the Kupffer's vesicle (KV) in phalloidin and DAPI stained Tg(*sox17:egfp*); *crb2a*-/- and sibling embryos at the 8-somite stage. **B**) Quantification of the number of KV cells in the embryos shown in (A). Each dot represents one embryo, black bars are median values, data analyzed by T-test, *** = P <0.001. **C**) *myl7* WISH staining in 36 hpf embryos injected with *crb2a* or control *tyr* sgRNAs to visualize heart looping. Fractions indicate the number of embryos with the phenotype shown over the total number of embryos analyzed for each condition. **D**) Quantification of laterality defects in *crb2a* crispant (shown in C) and *crb2a*-/- (not shown) embryos, data analyzed by Chi-Square test, **** = P <0.0001. **E**) *myl7* WISH staining in *sox32* morphants injected at the one-cell and 512-cell stages. Note that one-cell injected morphants have cardia bifida (2$^{nd}$ from left), whereas *sox32* morphants injected at the 512-cell stage exhibit laterality defects. Fractions indicate the number of embryos with the phenotype shown over the total number of embryos analyzed for each condition.

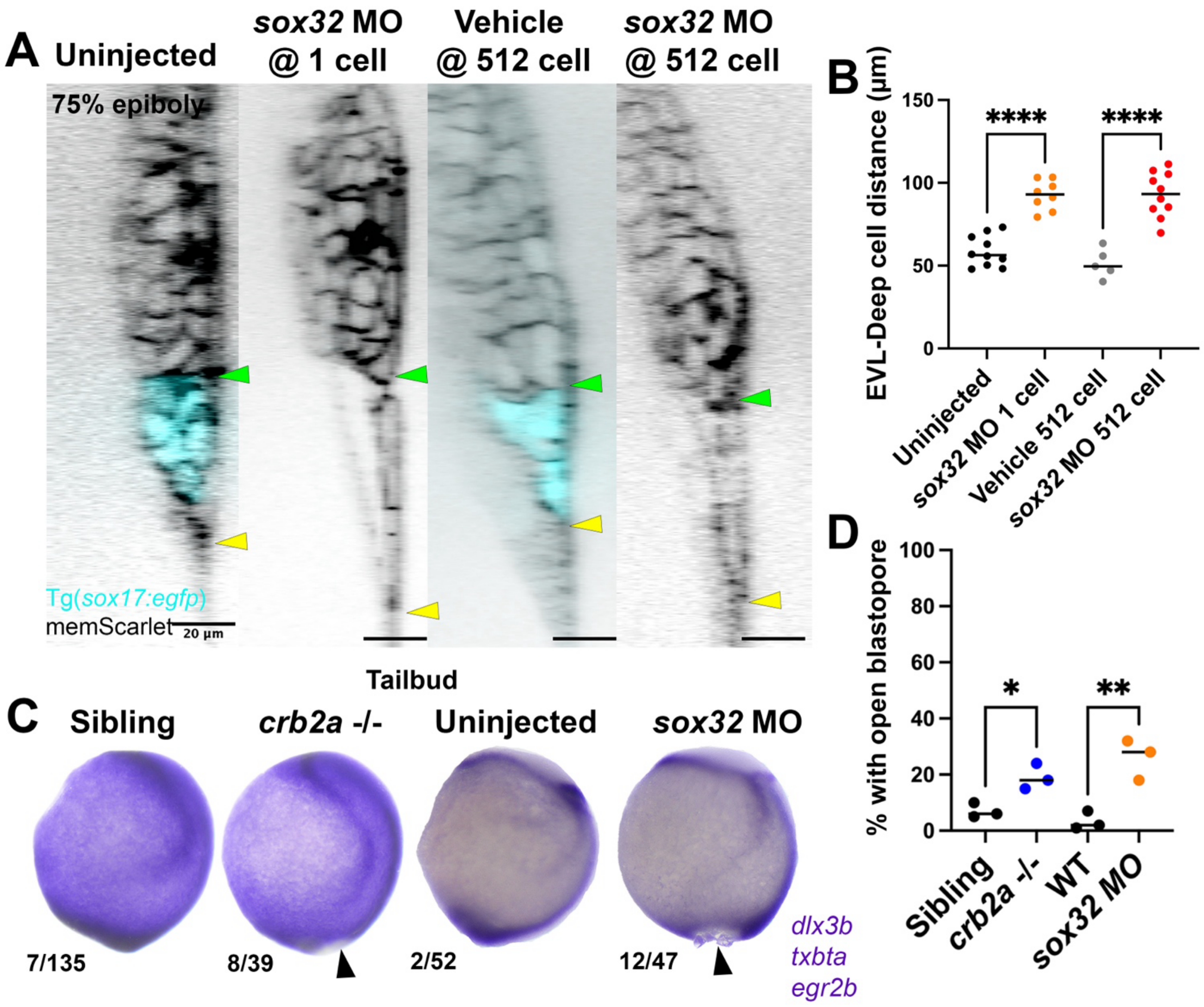


**Figure S4. Embryos with disrupted DFCs frequently have epiboly defects.** Related to Figures 3 and 4. **A**) Resliced optical sections of live Tg(*sox17:egfp*) *sox32* morphants at 75% epiboly, injected at the one-cell or 512-cell stages, and their respective controls showing decoupling of the enveloping layer (yellow arrowheads) and the deep cell layer (green arrowheads). **B**) Quantification of EVL-deep cell distance in the embryos shown in (A). Each dot represents a single embryo, Black bars are median values. Data were analyzed by T-test, **** = P <0.0001. **C)** Representative images of *crb2a*-/- mutants, *sox32* morphants, and sibling controls for the probes indicated at tailbud stage (10 hpf). Arrowheads indicate open blastopores. Fractions indicate the number of embryos per condition with open blastopores over the total number of embryos examined. **D**) Quantification of the % embryos of the conditions indicated with open blastopores. Each dot represents an independent clutch, black bars indicate median values. Data were analyzed by T-test, * = P <0.05, ** = P <0.01.

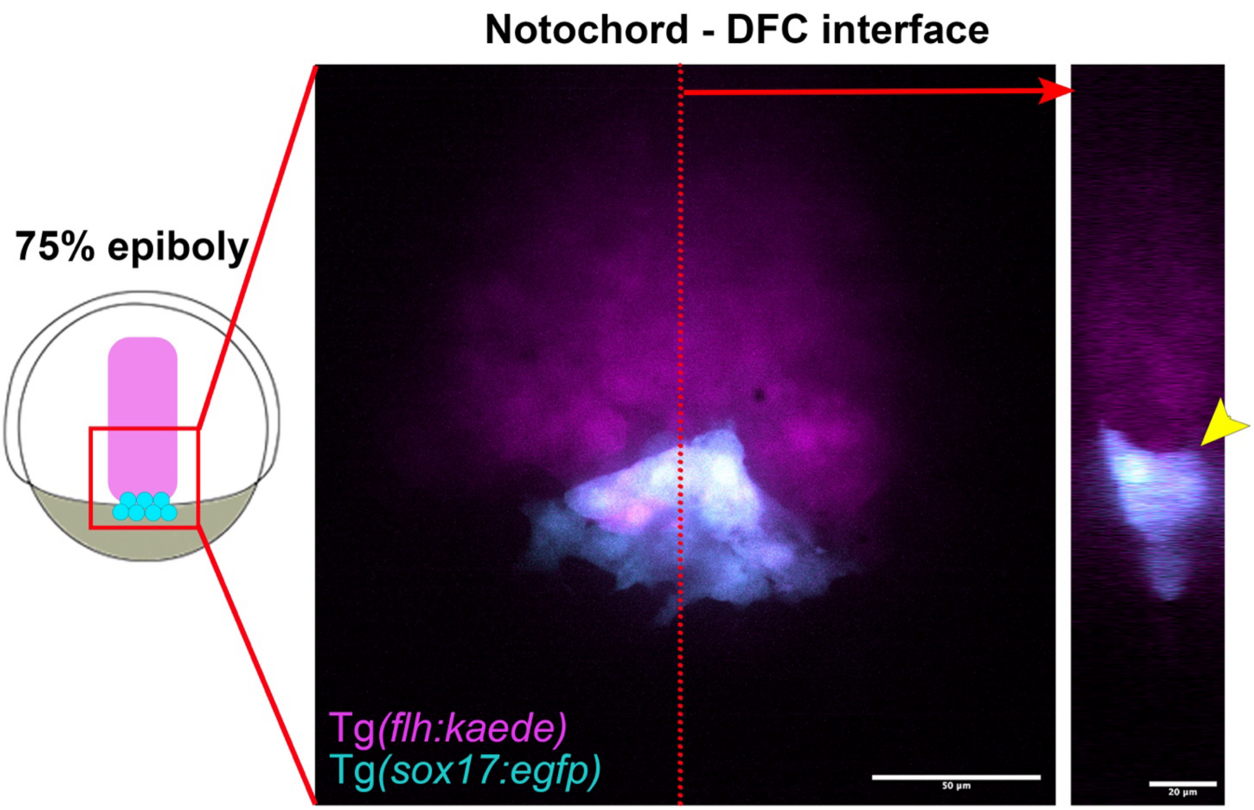


**Figure S5. DFCs are in direct contact with notochord cells during epiboly.** Related to Figure 5. 3-dimensional projection of DFCs (expressing *sox17:egfp*) and the developing notochord (expressing *flh:kaede*) in a WT embryo at 75% epiboly. Resliced optical YZ section (at the position of the red line) showing the saddle-like shape of the DFCs in contact with the posterior end of the notochord (yellow arrowhead).

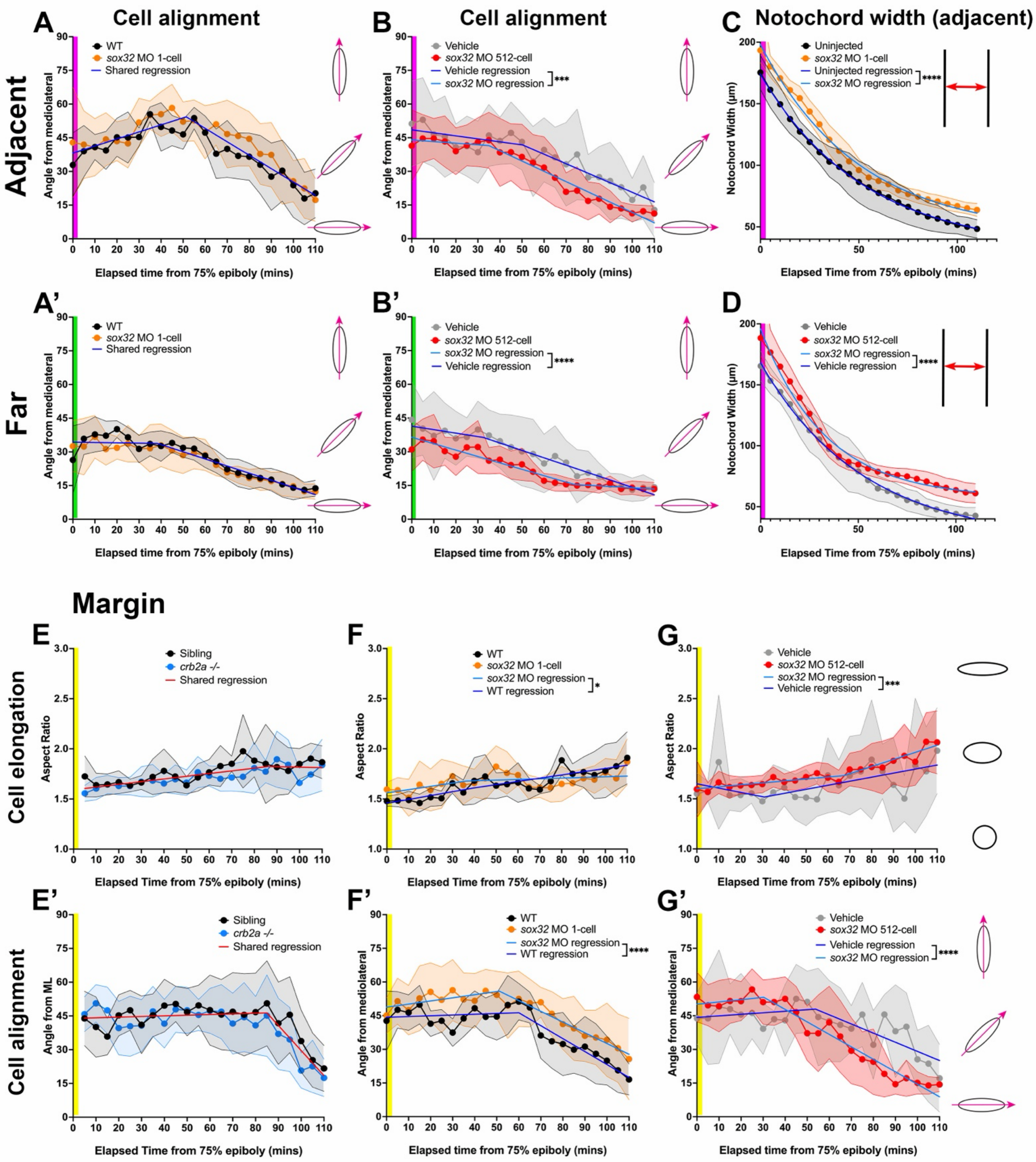


**Figure S6. Cell shape analysis of *crb2a-/-* mutants and *sox32* morphants during gastrulation.** Related to Figures 5 and 6. **A-B')** Quantification of cell alignment in the conditions and regions indicated. Graphs show medians and inter-quartile range. Data were fit with segmented lines using non-linear regression, differences were detected by sum-of-squares F-test, **** = P <0.0001. **C-D**) Quantification of notochord width over time in the adjacent region in embryos of the conditions indicated. Data were fit with a one-phase decay non-linear regression, differences were detected by sum-of-squares F-test, **** = P <0.0001. **E-G')** Quantification of cell elongation (top) and alignment (bottom) for cells in the margin region of embryos of the conditions indicated. Data were fit with segmented lines using non-linear regression, differences were detected by sum-of-squares F-test, **** = P <0.0001.

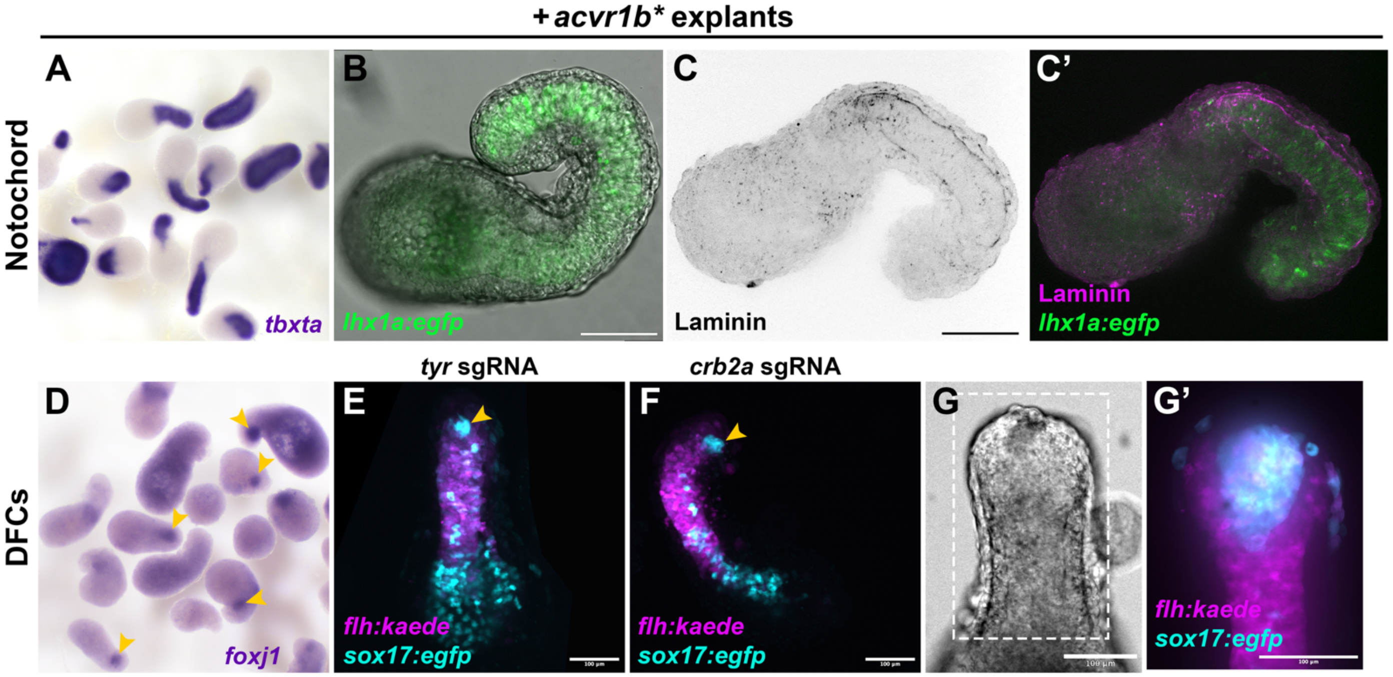


**Figure S7. Explants generate notochord-like structures and DFC-like cells.** Related to Figure 7. Representative images of zebrafish embryonic explants expressing *acvr1b** at 12 hpf (equivalent of 4-somite stage). **A-C')** Explants generate notochord-like structures expressing *tbxta* by WISH (A) and a mesoderm-specific *lhx1a:egfp* transgene (B, C'), and are surrounded by laminin detected by IF staining (C-C'). **D-F)** Embryonic explants produce cell clusters expressing DFC markers *foxj1* by WISH (D) and a *sox17:egfp* transgene (orange arrowheads) abutting *flh:kaede*-labeled notochord-like structures (E-F). **G')** Notochord boundaries can be observed by bright-field microscopy (G) and *flh:kaede* expression (G'). *sox17:egfp*+ DFC-like cells are located at the tip of the notochord-like structure (G').
